# Rigid granular scaffolds reveal transcriptional responses to cellular confinement

**DOI:** 10.64898/2026.11.21.768028

**Authors:** Shinny Sunny, Pei Ying Ng, Vaishnavi Rangaraj, Malhar Chitnis, Yixuan Li, Jia Wen Nicole Lee, Xu Gao, Hui Ting Ong, Paramasivam Kathirvel, Jin Zhu, Lindsay Riley, Tatiana Segura, Andrew W. Holle

**Affiliations:** Mechanobiology Institute, National University of Singapore, 117411, Singapore; Department of Biomedical Engineering, National University of Singapore, 117583, Singapore; Department of Biomedical Engineering, Duke University, Durham, NC, 27705, USA; Department of Medicine, Neurology, Dermatology, Duke University, Durham, NC, 27710, USA

## Abstract

A large variety of cells migrate through confined spaces *in vivo*, where mechanical confinement influences their fate and function. However, existing *in vitro* confinement models are low-throughput, limiting discovery of confinement-responsive genes and recovery of confined cells. Here, we present non-annealed glass microparticle scaffolds (GlaMS) as a scalable, high-throughput 3D confinement system with interconnected voids that mimic physiological confinement while enabling efficient cell retrieval. Using LOVAMAP, a computational tool that segments void spaces to estimate spatial constrictions, we quantified confinement levels in GlaMS and compared them with *ex vivo* interstitial spaces to identify the sphere size that best mimics *in vivo* confinement. Using human mesenchymal stem cells (hMSCs) as a mechanosensitive cell model, confinement within GlaMS was found to induce morphological changes, accelerated migration, altered YAP signaling, and enhanced osteogenic priming after replating, demonstrating transient confinement-induced effects. Taking advantage of the high throughput capabilities of GlaMS, we performed RNAseq on populations of confined cells and identified confinement degree-dependent shifts in gene expression. TMEM158 emerged as a previously unrecognized confinement-sensitive gene, was independently validated in microchannel assays, and was found by loss-of-function analysis to contribute to efficient confined migration. Together, this work establishes GlaMS as a scalable platform for population-level confinement mechanobiology, allowing recovery and molecular profiling of cells exposed to predictable three-dimensional confinement.

**Significance Statement:** Physical confinement is a key feature of tissue microenvironments, yet existing experimental systems are often too low-throughput for population-scale molecular analysis. We developed rigid granular scaffolds called “GlaMS” that generate predictable, physiologically relevant three-dimensional confinement while allowing efficient recovery of large cell populations. Using this platform, we show that confinement in GlaMS alters cell morphology, migration, mechanosensitive signaling, osteogenic priming, and global gene expression. Transcriptomic profiling identified TMEM158 as a previously unrecognized confinement-responsive gene, and functional studies showed that it supports migration through highly confined spaces. This work provides a scalable platform for studying population-level confinement mechanobiology, enabling discovery of molecular mechanisms that regulate cell behavior in mechanically confining environments.

## Introduction

Densely packed tissue environments in the body impose physical constraints on migratory cells as they traverse interstitial spaces. Intravital imaging has mapped these spaces to be approximately 3-30 µm in diameter^[1]^. Diverse cell types navigate such confining environments during key biological processes such as development^[2]^, immunosurveillance^[3,4]^, stem cell homing^[5–7]^ and cancer metastasis^[8]^. Navigating these spatial constraints require cells to undergo substantial morphological changes^[9,10]^, cytoskeletal remodelling^[11,12]^, nuclear deformation^[13–15]^, and activation of mechanosensitive signalling pathways^[16,17]^, which can collectively influence its gene expression and fate^[18]^. Given the prevalence of physical confinement *in vivo* and its potential to regulate cellular state, understanding how confinement influences cell behaviour and function is therefore essential.

Current confinement systems like PDMS microchannels, sandwich assays, and micropatterned substrates are engineered to recapitulate *in vivo* confinement and have been extensively used to understand how cells sense and respond to confinement. Even though these systems are quite useful for studying single-cell confined migration dynamics through imaging^[19]^, they are inherently low-throughput and difficult to scale. These platforms do not allow for the predictable and controlled confinement of large numbers of cells, nor do they enable easy extraction of confined cell populations for downstream applications. This low-throughputness limits discovery of new confinement sensitive genes using omic level analysis like bulk-RNA seq or use of confined cell populations for therapies. To address these issues, we propose granular scaffolds as a new approach for high throughput and high content confinement mechanobiology.

Granular scaffolds consist of microparticles that, when randomly close-packed in three dimensions, form interconnected void spaces, resulting in an inherently microporous structure. The size of interstitial void spaces can be tuned by adjusting the microparticle diameter. Microporous annealed particle (MAP) scaffolds are generally made from soft crosslinked hydrogel particles due to their tunable stiffness and injectability^[20–24]^. These have been applied in regenerative medicine to promote tissue repair during wound healing^[25–27]^ and to enhance vasculogenesis and stem cell infiltration when injected into brain tissue post-stroke^[28–30]^. Peptide-functionalized annealed microgels with macroporous void spaces have been engineered to recreate *in vitro* osteochondral interfaces^[31]^. MAP scaffolds have also been used to modulate immune cell behavior by controlling void space through particle size variation. For instance, macrophages cultured in MAPs with smaller void spaces exhibited a reduced inflammatory phenotype *in vitro*, whereas MAPs with larger pores implanted in mouse skin wounds facilitated greater immune cell infiltration and collagen regeneration^[32,33]^. However, the soft hydrogel microparticles used in these systems are deformable, allowing cell-generated forces to dynamically alter pore geometry and, consequently, the degree of confinement over time. This has been confirmed in the literature, as fibroblasts have been shown to deform soft polyacrylamide microparticles (∼6-14 kPa, ∼15 µm diameter)^[34]^. Recently, cells within a granular scaffold composed of soft polyacrylamide microparticles (0.5-5 kPa, 30-50 µm diameter) were found to deform individual microparticles, significantly changing their roundness^[35]^. In contrast, stiff glass-based microparticle systems would not be altered by cell-generated forces, allowing for the isolation of the effects of confinement without the confounding influence of matrix deformation. This distinction is critical for interpreting confinement-driven cellular responses and highlights the importance of considering particle stiffness when designing granular scaffolds for confinement studies. Granular scaffolds based on non-annealed stiff microsphere assemblies have previously been explored using silica beads to organize cellular systems^[36,37]^. Ordered hexagonal assemblies of silica beads have been used to create thin, cell-stabilized neuronal structures, where cells coat bead surfaces and form layered networks. However, these systems have been limited in their scalability and have not been utilized to explore confinement specifically. Overall, granular scaffolds have not yet been leveraged as a scalable, high-throughput tool for probing confinement effects in cells or for identifying confinement-responsive genes through transcriptomic analyses.

Here, we introduce Glass-based Microparticle Scaffolds (GlaMS), composed of micron-scale, non-annealed random close-packed glass beads with tight size distributions, as a robust and scalable platform for cell confinement. These stiff GlaMS resist deformation by cells and provide predictable, well-defined, and tunable void spaces that confine cells within a three-dimensional (3D) microenvironment mimicking *in vivo* dimensions. Because granular scaffolds can be increased in volume without altering void space dimensions, GlaMS provide a scalable route toward confinement of millions of cells while allowing for straightforward retrieval, an essential feature for translational applications.

Human bone marrow derived MSCs (hMSCs), was chosen as a mechanosensitive model to understand how granular confinement influences cell fate, function and transcriptional programs. Confinement of stem cells can significantly influence their differentiation, alter its migration behaviour, chromatin architecture, and modulate its proliferation^[11,16,18]^. In this study, MSCs were cultured within GlaMS for 3 or 7 days, after which they were either retrieved by trypsinization for immunofluorescence or lysed *in situ* for RNA sequencing. Because scaffold volume can be increased without altering local pore and constriction distributions, the architecture provides a route toward confinement of cell populations at the million-cell scale. We demonstrate that granular confinement leads to persistent reductions in cell and nuclear area post-extraction and replating, showcasing cells can retain transient confinement-induced effects which lasts up to 24 hours. Confined cells also migrate faster, activate YAP signaling, and show enhanced osteogenic priming. RNA sequencing confirms that GlaMS supports omics-scale analysis and reveals particle size-dependent transcriptional shifts. Furthermore, our system enabled the omics-assisted identification of confinement-responsive genes for the first time, revealing candidates that may play key roles in mediating stem cell adaptive responses to physical confinement. Altogether, this work positions granular scaffolds as a scalable, high-throughput platform for probing how confinement influences stem cell morphology, behavior, and fate.

## Results and Discussion

### Void spaces in GlaMS mimic in vivo confinement

Glass microparticle scaffolds provide several unique advantages for studying how physical confinement influences cell behavior (**Fig. 1A**). Traditional 3D confinement platforms such as bulk hydrogels or synthetic matrices limit downstream analysis, since recovery of encapsulated cells typically requires enzymatic digestion (e.g., collagenase), which is costly and can alter cell phenotypes including morphology, differentiation state, and migration properties. In contrast, densely packed non-annealed GlaMS containing tunable void spaces create a high-throughput confinement platform that enables long-term confinement of large cell populations, straightforward post-confinement recovery, and consequently, omics-scale analysis of these cell populations. GlaMS can also be adapted for confined migration assays, where cells seeded on top of the scaffold migrate through interstitial void spaces before collection of migratory cell population from the base. Together, these features make GlaMS a versatile platform for both mechanobiological studies and functional biological assays.

**Figure 1:**
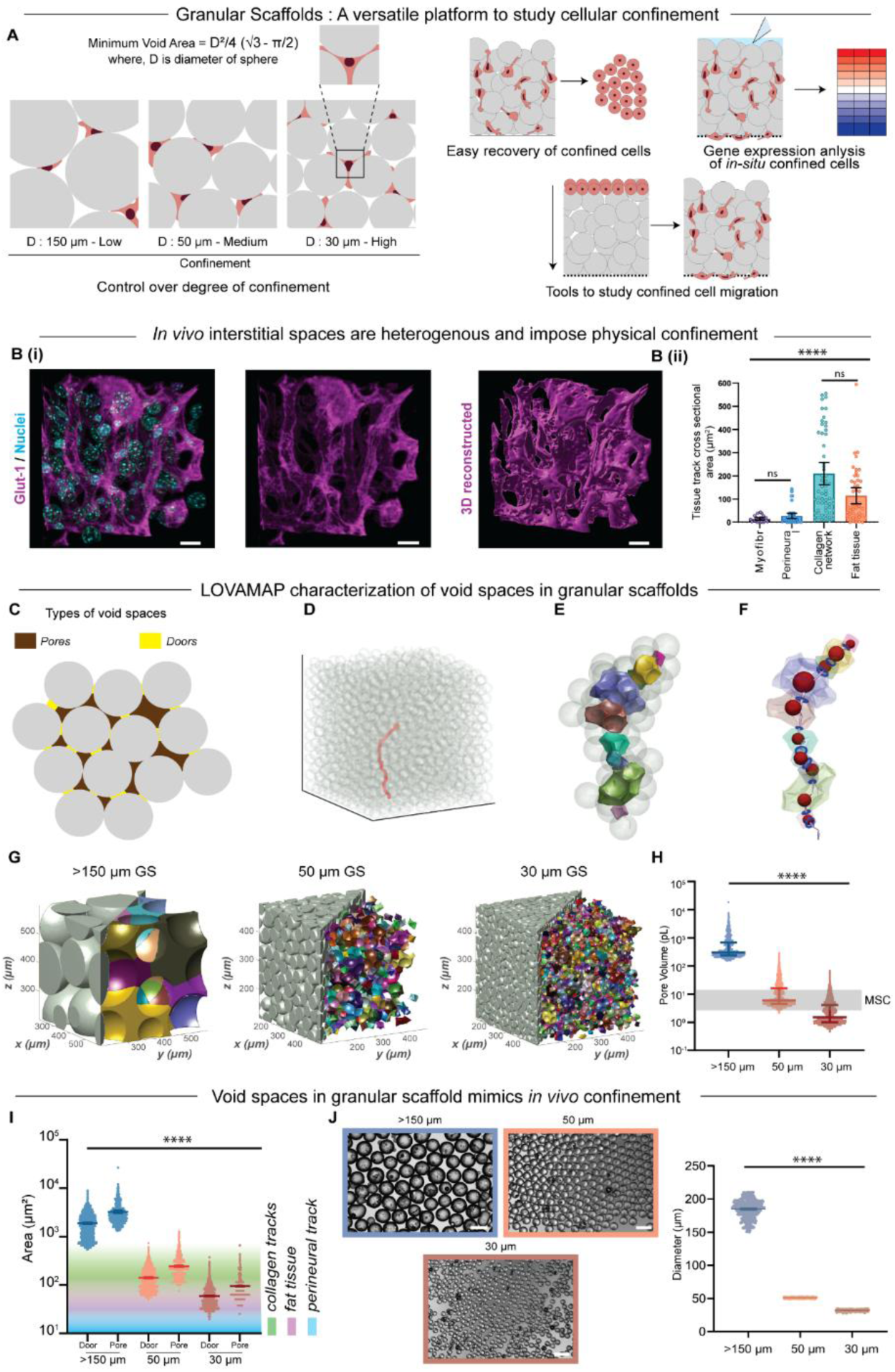
Engineering granular scaffolds for high-throughput cellular confinement. A) Schematic illustration of the applications of granular scaffolds in studying cellular confinement. Bi) Immunofluorescence images of mouse liver tissue slices stained for Glut 1 (purple) and nuclei (Hoechst) and 3D reconstructed in IMARIS to visualize interstitial spaces in vivo. Scale bar = 15 μm. Bii) Cross-sectional area of confining and heterogeneous in vivo spaces found in different tissues. Error bars represent 95% confidence intervals. C) Void spaces in granular structures can be divided into pores (open spaces) and doors (narrow regions). D) Simulated scaffold highlighting a single path (red) through the void space, generated by local void analysis using medial axis by particle configuration (LOVAMAP). Paths run from the center of scaffolds to the edge. Particles shown in transparent gray. E) Path from (D) highlighting 3D pores (in color). Surrounding particles are shown in transparent gray. F) Path from (D) highlighting largest enclosed sphere (red) of each pore, as well as doors between pores (blue circle). Pores shown in transparent color. G) Interior 3D pores (right half, in color) highlighting variations in pore size and shape between scaffolds packed with spheres of 150 μm, 50 μm, 30 μm diameters. H) Volume (in pL) of interior pores in granular scaffolds comprising each sphere population, as calculated by LOVAMAP. Error bars represents median with interquartile range. (30 µm diameter particles; N = 3 scaffolds), moderate confinement (50 µm diameter particles; N = 3 scaffolds), and unconfined (150 µm diameter particles; N = 10 scaffolds) I) Quantification of pore and door cross-sectional areas with error bar representing median with interquartile range. Underlay shows physiologically confining space ranges (by area) found in different tissues. J) Bright-field image of commercially sourced glass spheres with diameters of 150 μm, 50 μm, 30 μm, used to construct granular scaffolds. Scale bar = 100 μm. Frequency distribution graph of the glass sphere diameters. Error bars represent 95% confidence intervals.

A central feature of GlaMS is the tunability of void space architecture, allowing precise control over the degree of confinement. By adjusting bead diameter, void space size distributions can be scaled to match various biological contexts. Two key geometrical features of the void space have been defined^[23,38]^ within the packed structure: pores, or pockets of space enclosed by adjacent microparticles, and doors, which separate adjacent pores and exist within narrow passages (**Fig. 1C**). Together, these elements provide a framework for quantifying the spatial confinement experienced by cells. To systematically characterize pores and doors in 3D, we applied the Local Void Analysis using Medial Axis by Particle configuration (LOVAMAP) algorithm developed by Riley et al.^[23]^ This computational tool analyzes the void space of granular assemblies by first segmenting the space into 3D pores. Critically, LOVAMAP also identifies the location of doors, which allows for quantification of constriction points in a scaffold. These constrictions can hinder cell movement and are therefore important to consider when assessing cell migration paths and confinement degree during migration (**Fig. 1D-F, Fig. S1A-B**).

Using Houdini SideFX software, we simulated granular scaffolds comprising random close-packing of rigid spheres within a cubic container. The simulations modeled three size categories of GlaMS structures constructed from monodisperse glass particles with average diameters of 30 µm, 50 µm, and 150-210 µm (henceforth referred to as 150 µm), precisely matching the measured diameter distributions of different commercially available glass bead constituent materials. LOVAMAP analysis revealed distinct differences in pore and door size distributions across our three particle groups (**Fig. 1G**). Median pore volumes of approximately 1.5 pL, 5.94 pL, and 302.9 pL were calculated for GlaMS comprised of 30 µm, 50 µm, and 150 µm beads, respectively (**Fig. 1H**). As hMSCs typically have volume distributions ranging from 2.85 pL to 14.4 pL (corresponding to cell diameters of 15 - 30 µm), the pore spaces of 30 µm and 50 µm diameter scaffolds are generally smaller than a single cell and thus necessitate full 3D confinement. In contrast, the 150 µm GlaMS provides substantially larger void spaces. The minimum pore volume in this condition is approximately 153 pL, around 54-fold greater than the volume of a 15 µm diameter MSC **(Fig. 1H)**. Moreover, the minimum door cross-sectional area accessible to cells is approximately 537 µm², about threefold greater than the estimated cross-sectional area of a 15 µm diameter MSC (∼177 µm²). Together, this indicates that cells within 150 µm GlaMS can occupy pore spaces and migrate through doors without being confined. We therefore consider the 150 µm GlaMS condition as an unconfined 3D control.

To assess how well these engineered void spaces in GlaMS mimic native *in vivo* microenvironments, we extracted cross-sectional area dimensions of tissue tracks, such as adipose networks, perineural spaces, and collagen-dense regions, from published intravital imaging datasets^[1]^ (**Fig. 1Bii**). We then overlaid these dimensions onto LOVAMAP-derived pore and door cross-sectional size distributions (**Fig. 1I**). Notably, GlaMS composed of 30 µm and 50 µm glass spheres generated pore and door sizes that overlap with *in vivo* interstitial space dimensions, whereas the pore sizes in 150 µm scaffolds are supraphysiological and can thus act as an unconfined GlaMS experimental control. This validated our scaffold design for physiologically relevant confinement studies. Importantly, this comparison establishes similarity in the dimensions of spatial confinement found in GlaMS as compared to *in vivo* spaces, rather than suggesting that GlaMS reproduces the complete mechanical and biochemical properties of native tissues. Moreover, the glass particles are effectively non-deformable by cellular forces preventing any changes to the confinement during cell migration, allowing us to decouple effects of spatial confinement from matrix remodelling.

We further correlated our scaffold findings to physiological conditions by imaging interstitial spaces in murine tissues. Mouse quadriceps muscle was sectioned, optically cleared, and stained for collagen, actin, and nuclei (**Fig. S2A**). In these samples, we observed deformed nuclei, indicating that cells within interstitial compartments experience confinement from the surrounding extracellular matrix. Mouse liver tissue was also stained for Glut1, a ubiquitous plasma-membrane glucose transporter that mediates glucose uptake and homeostasis in mammalian cells^[39]^. By labeling cell membranes and nuclei and digitally inverting the resulting images, we could visualize the interstitial spaces *in situ* (**Fig. 1Bi**, **Fig. S2B**, **Supplementary Video 1**). These interstitial spaces measured ∼15 µm across, representing the mechanical constraints cells experience *in vivo*. Importantly, the alternating pore and door architecture of GlaMS void space captures the heterogeneity of irregular interstitial spaces found *in vivo* as seen in our visualization of mouse tissue (**Fig. 1Bi**). Based on these comparisons, we utilized three bead diameters: 30 µm (extreme confinement), 50 µm (moderate confinement), and 150 µm (unconfined). To fabricate GlaMS, we used commercially available monodisperse glass microparticles in the specified sizes, which settle under gravity in a random close-packed manner (**Fig. 1J**).

The smallest pore in a random close-packing of monodispersed rigid spheres is the space enclosed by four equal-sized spheres touching each other^[38,40]^. The doors exiting this pore represent the narrowest circular passages that cells encounter, and their area can be calculated as shown in **Fig. 1A**. Alternatively, the minimum void area a cell has access to when migrating through this region can be represented by the non-particle triangular space enclosed by three particles. The 3D pore and door sizes in GlaMS scale proportionally with particle diameter, as evidenced by the larger void volumes found in scaffolds packed with larger spheres (**Fig. 1H**).

### GlaMS enable long-term, high-throughput cellular confinement

To investigate the mechanosensitive effects of cellular confinement with GlaMS, 30 µm, 50 µm, and 150 µm glass microparticles were coated with collagen and dispersed as a monolayer into low-attachment 6-well plates. Low-attachment plates ensured cell attachment to the microparticle surface rather than the plate surface. hMSCs were then seeded onto the particles and allowed to attach for 24 hours at 37 °C. Cell-laden microparticles were then transferred into custom 24-well transwell inserts, allowing the particles to spontaneously settle into a random close-packed arrangement, confining the cells within the 3D void spaces (**Fig. S7**). The use of transwell inserts facilitated routine media exchange without disrupting the structural integrity of the granular scaffolds, thereby supporting sustained confinement over extended culture periods. Collagen coating of glass particles allowed improved pre-confinement cell adhesion on the microparticles, and higher post-confinement cell yield during recovery (**Fig. S3**). During initial seeding of cells onto the microparticle monolayer, they attach and wrap around the bead surface (**Fig. S5C**). Upon 3D packing, cells are able to spread to other bead surfaces through surrounding doors and pores (**Fig. 2D, Fig. S6A**).

**Figure 2:**
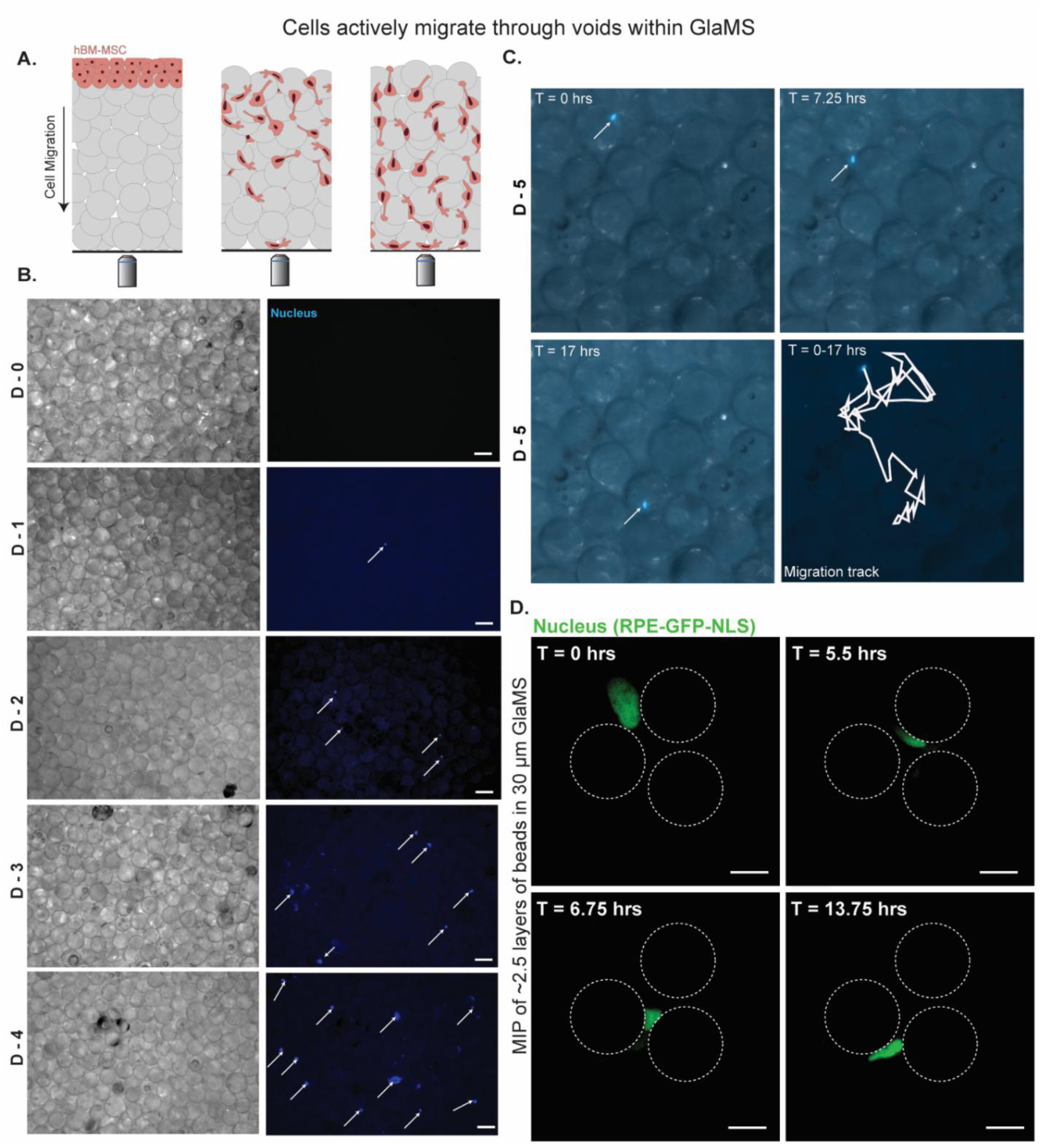
Cells undergo active migration through interstitial void spaces in GlaMS. (A) Schematic illustrating cell seeding at the upper surface of GlaMS and subsequent active migration through interconnected pores and doors formed between packed beads.(B) Brightfield and fluorescence images of hMSCs over time (Day 0 - Day 4) within 30 µm GlaMS. Scale bars: 30 µm (C) Time-lapse imaging of a representative cell migrating over 17 h. Arrows indicate cell position at different time points. (D) High-resolution confocal imaging of nuclear deformation during migration through confined pores between beads. RPE nuclei (RPE-GFP-NLS, green) deform as they migrate through the narrow interstitial “doors” between adjacent beads (outlined by dashed circles). Images represent maximum intensity projections (MIP) over ∼2.5 bead layers in 30 µm GlaMS. Scale bars: 20 µm.

Conventional microchannel-based confinement systems typically allow for the collection of roughly 100 cells post-confinement, significantly limiting scalability^[18]^. Due to the 3D nature of GlaMS, it was possible to recover over 15,000 cells from a single well after seven days of the most extreme confinement using a convenient harvesting protocol (**Fig. S4B**). This increase in throughput comes at the cost of confinement homogeneity, as microchannel assays guarantee that cells experience specific confinement widths, although it is important to note that the GlaMS present a distribution of void sizes that is well matched to the range of interstitial confinement found *in vivo* (**Fig. 1I**). Although our current experiment demonstrates recovery of over 15,000 cells from a single well of the most confining GlaMS, the scaffold itself is scalable **(Fig. S15)** without any changes to the local door-pore architecture. Thus, the current study experimentally established increased throughput relative to conventional microchannel systems while providing a framework that could be expanded further when larger cells numbers are required.

For confinement within a granular scaffold to be meaningful, the majority of cells must not be able to move the microparticles and remodel the 3D network, as this would result in an alteration of the distribution of confining pore volumes and door areas. Thus, a fully jammed microparticle network is a requirement. To assess how much of the scaffold maintains this jammed configuration and provides consistent confinement, we performed a simple bead displacement assay. Increasing layers of 30 µm glass particles (the smallest particles used in this study) were stacked onto a monolayer of adherent cells, and bead movement at the bottom layer was imaged over time. GlaMS with over 11 layers of particles resulted in a fully jammed state at the bottom layer, while cell-induced microparticle movement was observed in GlaMS with less than 9 layers (**Fig. S4A**, **Supplementary Videos 2-3**). As all the confining GlaMS used in subsequent experiments were prepared with over 100 layers, we conclude that at least 80% of encapsulated cells experience consistent confinement ranges.

### Cells undergo active migration through void spaces in GlaMS

Initial plating of cells on a monolayer of beads ensures even distribution of cells throughout the scaffold, but it is unknown if cells migrate through the microparticle void space network or simply remain stationary within the pores. Given the refractive index mismatch between the glass microparticles (RI ∼1.5) and their surrounding media (RI ∼1.33), direct visualization of cells deep within the GlaMS is not feasible (**Fig. S5A**), even when higher refractive index solutions were used (**Fig. S5B**). However, using a silicone immersion objective with a long working distance and relatively high numerical aperture in conjunction with the far-red dye SiR-DNA (as longer wavelengths can reduce scattering), we were able to image 2-3 layers (50-75 µm) deep into the GlaMS, providing representative visualization of cell migration within the scaffold (**Fig. S6A**). Time-lapse microscopy of cells within the GlaMS confirmed that stem cells do migrate within the scaffold, that their nuclei are significantly deformed during this process, and that the fully jammed microspheres do not move or deform as cells migrate between them (**Fig. 2D**).

To strengthen our observations that long-distance confined migration occurs in the scaffold, we plated cells at the top of the 30 µm GlaMS scaffold prior to time-lapse microscopy focused at the bottom surface of the scaffold (**Fig. 2A**). The first cells were observed arriving at the bottom surface (∼1,000 µm below the upper surface) roughly 24 hours after seeding, which is consistent with the approximate cell migration speed found in microchannel confinement (∼0.7 µm/min) (**Fig. 2B**). No cells were observed at the bottom surface immediately after seeding, proving that cells do not passively fall through the void spaces to the bottom of the well. We also confirmed that the cells arriving at the bottom are viable via time-lapse microscopy that showed continued cell migration (**Fig. 2C**), with roughly 85% of all cells found to be motile (**Fig. S6B**).

### Granular confinement causes altered cell morphology and migration dynamics

To examine the effects of granular confinement on cell morphology, hMSCs were subjected to 3 days of confinement in GlaMS constructed using 30 µm, 50 µm, and 150 µm particles. As imaging throughout the GlaMS column is challenging due to the long working distance required, we transferred cells out of the scaffold and onto collagen-coated coverslips for downstream analysis of confinement-induced cellular responses **(Fig. S8)**. Cells were gently released by trypsinization and replated for 3 hours (long enough to allow for cell spreading but short enough to prevent loss of confinement-induced phenotype changes) before fixation and staining for actin and nucleus (**Fig. 3A**).

**Figure 3:**
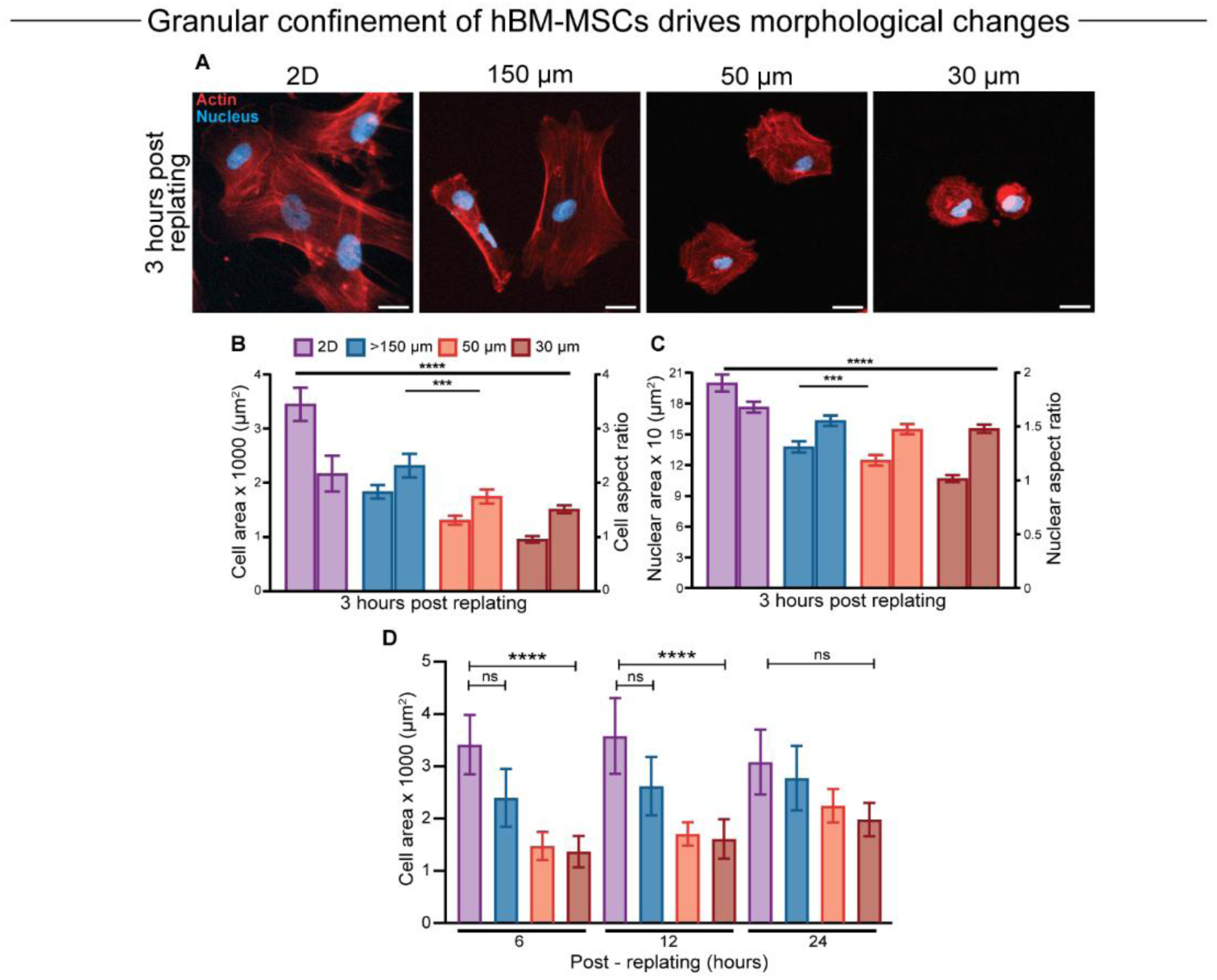
Granular confinement of stem cells drives morphological changes. A) Immunofluorescence images of hMSCs confined in granular scaffolds for 3 days, extracted, and replated for 3 hours before fixation and staining for actin (red) and nuclei (blue). Scale bar = 20 µm. B) Quantification of cell area and cell aspect ratio. N = 3, n = 306. C) Quantification of nuclear area and nuclear aspect ratio. N = 3, n = 338. D) Cell area quantified for cells confined in GlaMS and replated for 6, 12, 24 hours. N = 2, n = 30 (6 hours), 30 (12 hours), 50 (24 hours). All graphs have error bars represented as 95% confidence interval. Kruskal-Wallis test with Dunn’s multiple comparison test for statistical significance. *P <= 0.05, **P <= 0.01, ***P <= 0.001, ****P <=0.0001.

Cells within GlaMS are exposed to three key mechanical cues: (i) substrate stiffness, (ii) curvature of the glass microspheres, and (iii) confinement arising from bead packing. To isolate the effect of confinement from substrate stiffness, glass 2D controls were used to ensure that stiffness was consistent across conditions and does not account for the observed phenotypic differences. To further exclude the effects of curvature, we also employed a 2.5D control in which cells were seeded on glass microspheres of the same diameter (30 µm, 50 µm and 150 µm) but were not jammed (**Fig. S9A**). In this setup, cells experience bead curvature without confinement.

Quantification of cellular and nuclear spread area post-replating revealed a significant reduction in both metrics and enhanced nuclear rounding compared to unconfined 2D controls (**Fig. 3B-C**), with the most pronounced area reduction and nuclear rounding observed in cells previously confined in the highest confinement condition (30 µm GlaMS). This reduction in cell and nuclear area is proportional to the degree of prior granular confinement. While a reduction in cell and nuclear area in the 2.5D condition was observed compared to 2D **(Fig. S9B-C)**, the reduction in the 3D GlaMS is significantly greater, suggesting that differences in cell phenotype mainly arise from confinement within the packed bead structure, rather than from substrate stiffness or curvature. As seen in other confinement platforms, confining microenvironments physically limit cell spreading, which is reflected as smaller projected cell and nuclear areas^[15,18]^. The fact that this change in cell and nuclear spread area was still present 3 hours post-replating suggests that these morphology changes are not solely due to the ongoing presence of confinement, but at least partially due to a cellular adaptation to confining spaces that persists for some period of time after the cell enters a new environment. To determine how long these changes last, we cultured hMSCs in GlaMS for 3 days and then replated them for 6, 12, and 24 hours. Confined cells retained reduced cellular areas compared to unconfined controls after 6 and 12 hours, but this difference was lost 24 hours post-confinement (**Fig. 3D**).

Like other adherent cells, hMSCs have been found to migrate faster in confinement, a phenomenon that has been attributed to the mesenchymal-to-amoeboid transition of migration mode^[15,18,41]^. To test whether cells can retain this increased migratory capacity in a new environment, we confined hMSCs in GlaMS for 3 days, then performed time-lapse microscopy one hour after replating. At 6 hours post-replating, cells from 30 µm GlaMS confinement migrated significantly faster than those that were exposed to 50 µm or 150 µm scaffolds (**Fig. 4G**). However, like the changes in spread area, this difference was lost by 24 hours, suggesting that cell morphology and migration dynamics induced by confinement can be retained for limited periods of time post-replating.

**Figure 4:**
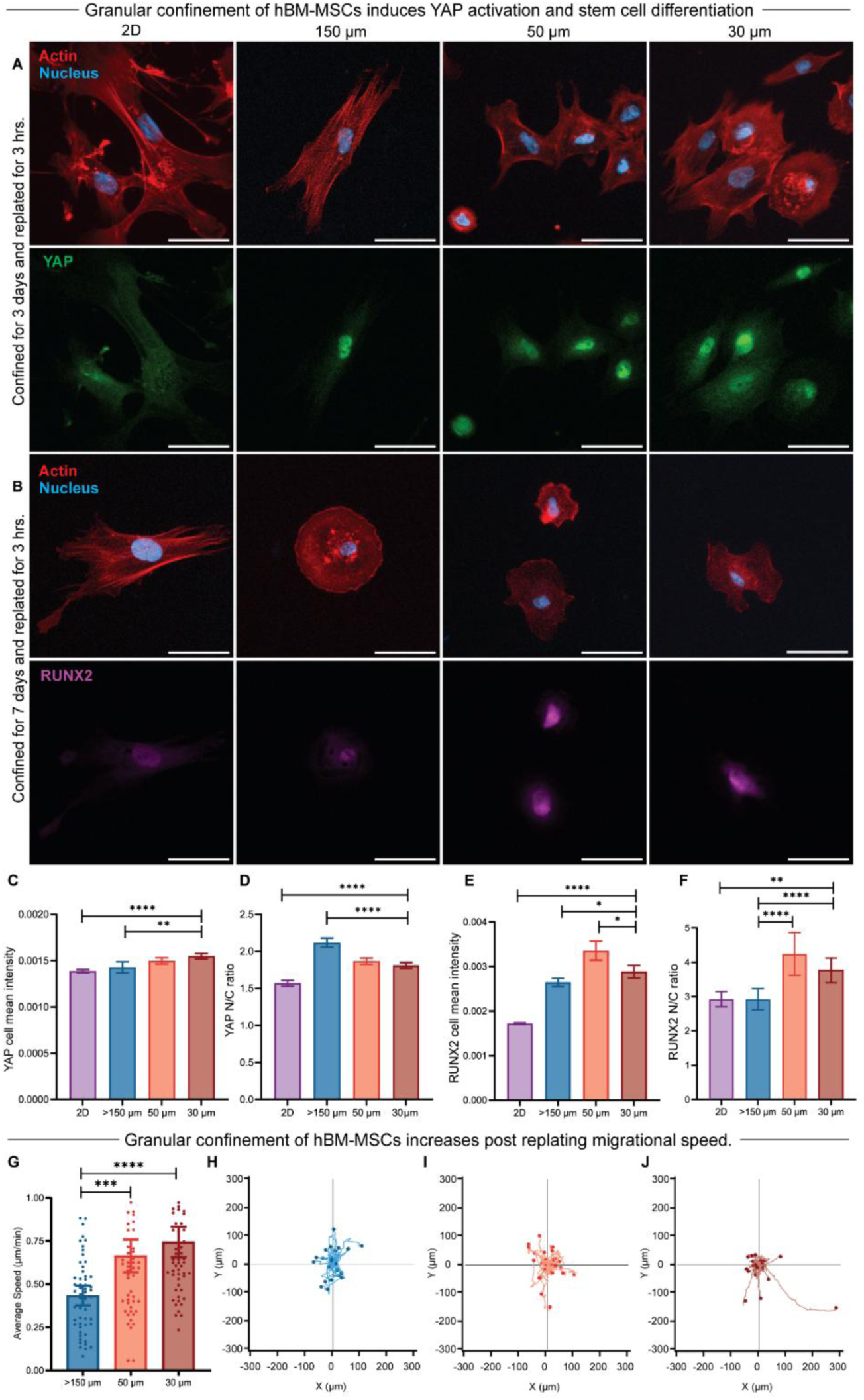
Granular confinement regulates stem cell fate and behavior. A) hMSCs were confined in a granular scaffold for 3 days, then replated for 3 hours on collagen-coated glass coverslips and stained for actin (red), nuclei (blue), and YAP (green). Scale bar = 20 µm. B) hMSCs were confined in granular scaffolds for 7 days, then replated for 3 hours on collagen-coated glass coverslips and stained for actin (red), nuclei (blue), and RUNX2 (purple). Scale bar = 50 µm. C-D) Quantification of the YAP cell mean intensity and N/C ratio in the confined cells. N = 3, n = 197, 307, 338, 339, respectively. E-F) Quantification of the RUNX2 cell mean intensity and N/C ratio in the confined cells. N = 3, n = 416, 464, 239, 229, respectively. G) Quantification of average migration speed. hMSCs were confined for 3 days, then replated on collagen-coated glass coverslips and live imaged to track cell migration over 6 hours. N = 2, n = 55, 55, 58, respectively. All graphs have error bars represented as 95% confidence interval. Kruskal-Wallis test with Dunn’s multiple comparison test for statistical significance. *P <= 0.05, **P <= 0.01, ***P <= 0.001, ****P <=0.0001. H-J) Trajectory rose plots showing the migration paths of hMSCs after replating on collagen-coated glass coverslips following 3 days of confinement in GlaMS.

### Granular confinement alters mechanosensitive signalling and stem cell fate markers

YAP, a mechanosensitive transcriptional co-activator and central effector of the Hippo signaling pathway, is known to translate mechanical cues into a variety of gene expression programs, including those controlling stem cell differentiation^[42,43]^. Nuclear deformation is a key trigger for YAP activation in hMSCs, as observed by elevated YAP nuclear localization in cells migrating through confining microchannels, a mechanical cue that subsequently directs their cell fate^[18]^. This led us to hypothesize that granular confinement, which similarly imposes physiological confinement and deforms the nucleus **(Fig. 2D, Fig. S6A)** would activate the YAP pathway in hMSCs and influence cell fate decisions.

hMSCs were confined in GlaMS for 3 days and replated for 3 hours before fixing and staining for YAP. Cells confined in 30 µm GlaMS, which experience the greatest degree of nuclear deformation, displayed significantly higher total cellular YAP expression after replating compared to unconfined cells or those in 150 µm GlaMS **(Fig. 4A,C)**. Quantification of the ratio of nuclear YAP to cytoplasmic YAP after replating revealed the opposite trend, with higher translocation observed in cells from 150 µm GlaMS compared to those in 30 µm GlaMS **(Fig. 4D)**. Nuclear YAP intensity showed no significant change as a function of scaffold **(Fig. S10).** As YAP nuclear translocation is a dynamic and often short-lived process^[44]^ that is strongly influenced by cell spread area^[42]^, it is difficult to fully decouple the effects of post-replating cell size from those of prior granular confinement. However, the overall increase in total YAP expression in highly confined hMSCs supports the idea that confinement alone, decoupled from substrate stiffness, can act as a mechanical cue to modulate mechanosensitive YAP signaling. This is particularly notable because most granular scaffold studies have examined YAP activation in the context of varying stiffness, where increases in microparticle stiffness and size promote enhanced cell spreading and are associated with YAP nuclear translocation^[45,46]^.

As altered YAP signalling has been shown to play a role in stem cell lineage specification^[42,47]^, we next investigated whether granular confinement can influence hMSC differentiation. Cells were confined in GlaMS for 7 days and replated for 3 hours before fixing and staining for RUNX2, an early osteogenic transcription factor. hMSCs in 50 µm and 30 µm GlaMS displayed significantly elevated RUNX2 expression, with increases in both total cellular mean intensity and nuclear translocation (N/C ratio), compared to 150 µm and unconfined 2D controls **(Fig. 4B,E,F**), consistent with previous confinement studies^[18,48]^. Interestingly, RUNX2 expression was higher in 50 µm GlaMS than in the 30 µm GlaMS confinement, suggesting that there may be an optimal degree of confinement for osteogenic priming, rather than a linear increase with the degree of confinement, with 50 µm GlaMS providing a more favorable mechanical environment than 30 µm, consistent with previous studies examining stiffness-dependent behavior of stem cells on 2D substrates^[49]^. The observed increase in RUNX2 nuclear localization in the absence of any osteogenic induction media indicates that granular confinement alone is sufficient to prime hMSCs toward osteogenesis.

### Identification of novel granular confinement-responsive genes via global transcriptomic profiling

Despite growing interest in confinement mechanobiology, a comprehensive transcriptional profile of cells confined within physiologically relevant dimensions has not been generated. Such knowledge is critical for understanding how confined stem cells behave during *in vivo* migration and how their functions or fates can be regulated *in vitro* and ultimately in therapeutic settings. Leveraging the high-throughput nature of our GlaMS platform, we performed a transcriptomic analysis that would not be feasible using conventional confinement systems. To this end, hMSCs were confined in 30 μm, 50 μm, and 150 μm GlaMS for 7 days. Importantly, unlike prior experiments that involved cell extraction and replating, cells were lysed directly within the scaffold using TRIzol to allow collection of expression signatures of *in situ* confined cells **(Fig. S11A)**. RNA-seq was performed on the extracted lysates, resulting in the identification and quantification of over 15,000 genes.

Principal component analysis (PCA) revealed clear transcriptional segregation **(Fig. S11B)**. PC1, accounting for 53% of the variance, sharply distinguished the 2D unconfined controls from the 3D GlaMS samples. PC2 (25%) separated the GlaMS conditions by confinement degree, with 150 μm samples clustering separately from the 30 μm and 50 μm conditions. Although cells in both 150 μm GlaMS and on 2D glass coverslips are unconfined, they remain transcriptionally distinct. To determine how the degree of confinement within GlaMS influences cell function, we identified differentially expressed genes (DEGs) for each pairwise comparison using a stringent threshold of adjusted p ≤ 0.05 and fold change ≥ 2. High confinement in 30 µm GlaMS, compared with the unconfined 150 µm GlaMS condition, resulted in 647 DEGs, indicating a strong transcriptional response to high spatial confinement **(Fig. 5A)**. In contrast, moderate confinement in 50 µm GlaMS, compared with 150 µm GlaMS, resulted in fewer DEGs (331 genes), suggesting a milder molecular shift **(Fig. 5B)**. The lowest number of DEGs were observed between high confinement 30 µm GlaMS vs the moderate confinement 50 µm GlaMS (32 genes), indicating that these two confined conditions elicited highly similar transcriptional responses **(Fig. 5C).** Comparison with the 2D unconfined control revealed a larger number of transcriptional differences, with 1,724 DEGs in 30 µm GlaMS versus 2D and 1,094 DEGs in 50 µm GlaMS versus 2D **(Fig. 5D-E)**. Importantly, even the unconfined 150 µm GlaMS condition differed substantially from the unconfined 2D control, with 1,811 DEGs identified between these conditions **(Fig. 5F)**. Consistent with these pairwise comparisons, heatmap visualization using row-scaled z-scores and hierarchical clustering showed broadly opposing expression patterns between 2D and 3D GlaMS conditions **(Fig. 5G)**. The 30 µm and 50 µm confinement conditions clustered closely together and exhibited highly similar gene-expression profiles, whereas the unconfined 150 µm GlaMS condition formed a distinct group. Together, these findings support a confinement-associated transcriptional shift within the 3D GlaMS environment. To specifically identify confinement-responsive genes while minimizing transcriptional differences arising from dimensionality, subsequent analyses therefore focused on comparisons of DEGs between confined 30 and 50 µm GlaMS and the unconfined 150 µm GlaMS. Gene Set Enrichment Analysis (GSEA) showed that pathways related to nucleocytoplasmic transport and nuclear envelope organization were significantly upregulated in highly confined 30 μm cells **(Fig. 5H)**, supporting the notion that confinement-induced nuclear deformation can actively regulate nuclear import/export dynamics^[50]^. In addition, genes involved in the negative regulation of locomotion were downregulated in 30 µm confined cells, in line with our earlier observations of increased migration of highly confined stem cells.

**Figure 5:**
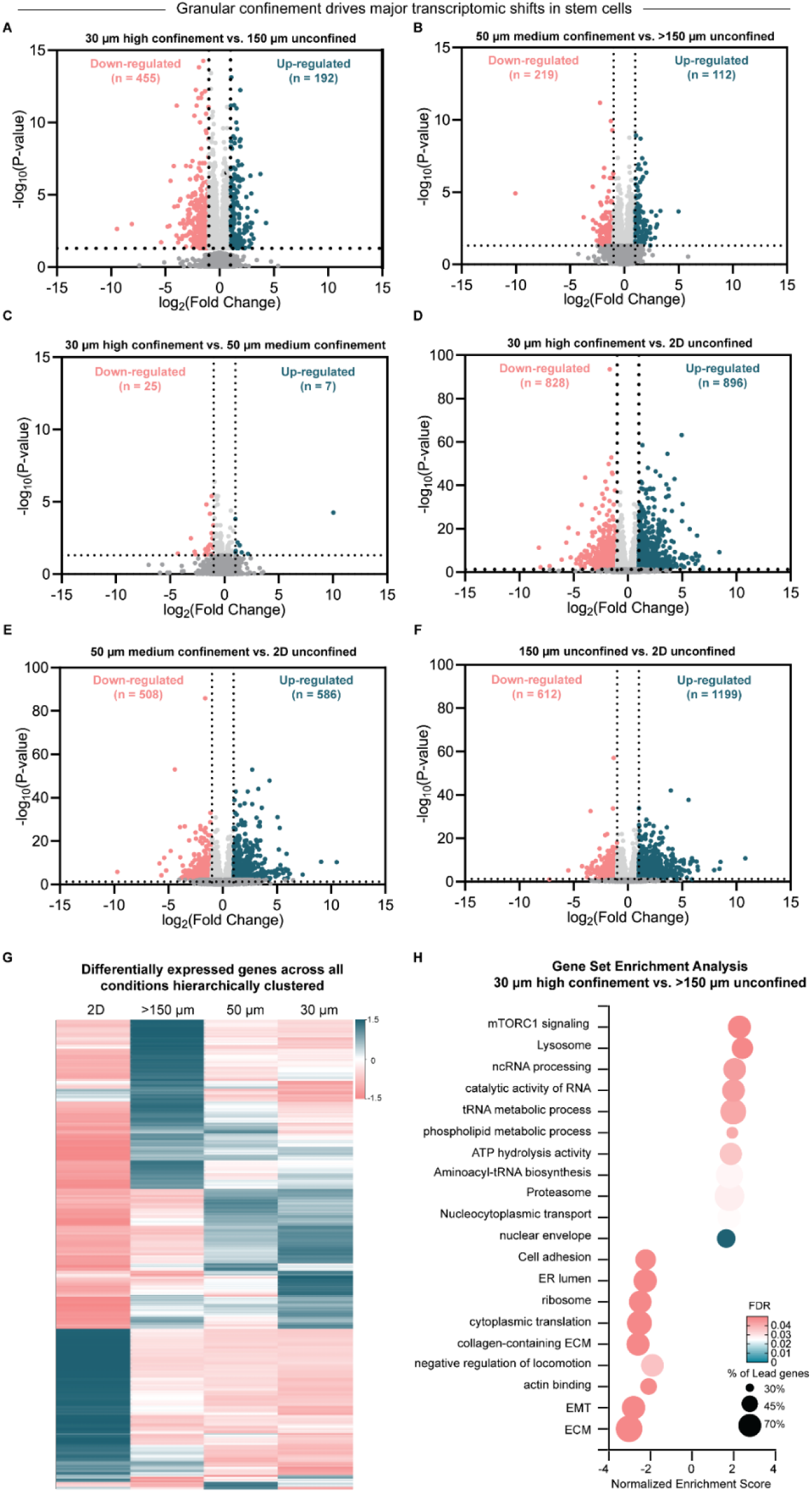
Transcriptomic analysis of stem cells following 7-day confinement in granular scaffolds. A) Differentially expressed genes in 30 µm GlaMS vs 150 µm GlaMS. B) Differentially expressed genes in 50 µm GlaMS vs 150 µm GlaMS. C) Differentially expressed genes in 30 µm GlaMS vs 50 µm GlaMS. D) Differentially expressed genes in 30 µm GlaMS vs 2D. E) Differentially expressed genes in 50 µm GlaMS vs 2D. F) Differentially expressed genes in 150 µm GlaMS vs 2D. All volcano plots have the x-axis representing log2fold change, and the y-axis showing - log10 adjusted p-values. Significantly upregulated genes (log2FC > 1, adjusted p < 0.05) are shown in green, while significantly downregulated genes (log2FC < -1, adjusted p < 0.05) are shown in pink. G) Heatmap showing differentially expressed genes across all conditions. Rows represent genes, columns represent experimental conditions. Gene expression values are normalized with Z-score, with hierarchical clustering applied. H) Bubble plot of enriched pathways among differentially expressed genes between 30 µm and 150 µm GlaMS conditions. Bubble size corresponds to the number of genes associated with each pathway, and color intensity indicates statistical significance. Pathway enrichment was performed using GSEA.

As transcriptomic approaches have never been applied to cells under varying levels of physiologically relevant confinement, we next sought to identify confinement-sensitive genes. Analogous to light switches that can be either flipped on or off and dimmer knobs that provide gradual changes in brightness, we classified confinement-responsive genes into “Switch” and “Dimmer” categories. “Switch” genes were defined as genes that exhibited a discrete transcriptional change at a specific level of confinement. These were further divided into two types. Type 1 Switch genes **(Fig. 6A)** were significantly up- or downregulated in 30 µm versus 150 µm and in 30 µm versus 50 µm GlaMS but were not significantly altered between 50 µm and 150 µm GlaMS. These genes therefore exhibited a marked transcriptional change only under high or extreme confinement. Type 2 Switch genes **(Fig. 6C)** were significantly altered in both 30 µm versus 150 µm and 50 µm versus 150 µm comparisons but were not significantly different between 30 µm and 50 µm GlaMS. These genes therefore responded similarly to both moderate and extreme confinement, representing a confinement-onset response in which gene expression changes once a threshold level of confinement is reached. In contrast, “Dimmer” genes **(Fig. 6B)** are genes that exhibited a gradual but consistent increase or decrease in expression from 150 µm to 50 µm to 30 µm. These genes were significantly different in all three pairwise comparisons, 30 µm versus 150 µm, 50 µm versus 150 µm, and 30 µm versus 50 µm, and exhibited a confinement dose-dependent change in expression. In all these cases, genes were classified as significantly differentially expressed at adjusted *p* ≤ 0.05 and further grouped by the magnitude and direction of log_2_FC (see Methods). Mechanosensitive “*Switch*” and “*Dimmer*” behavior has been previously observed on 2D substrates, where lineage-specific differentiation markers have been shown to peak exclusively at defined stiffness levels while certain mechanosensitive proteins like Lamin A show gradual, dose-dependent expression^[49]^. After selecting the DEGs that fulfill these criteria, we converted their transcript-per-million (TPM) values to Z-scores across all confinement conditions, enabling direct comparison of relative expression levels.

**Figure 6.**
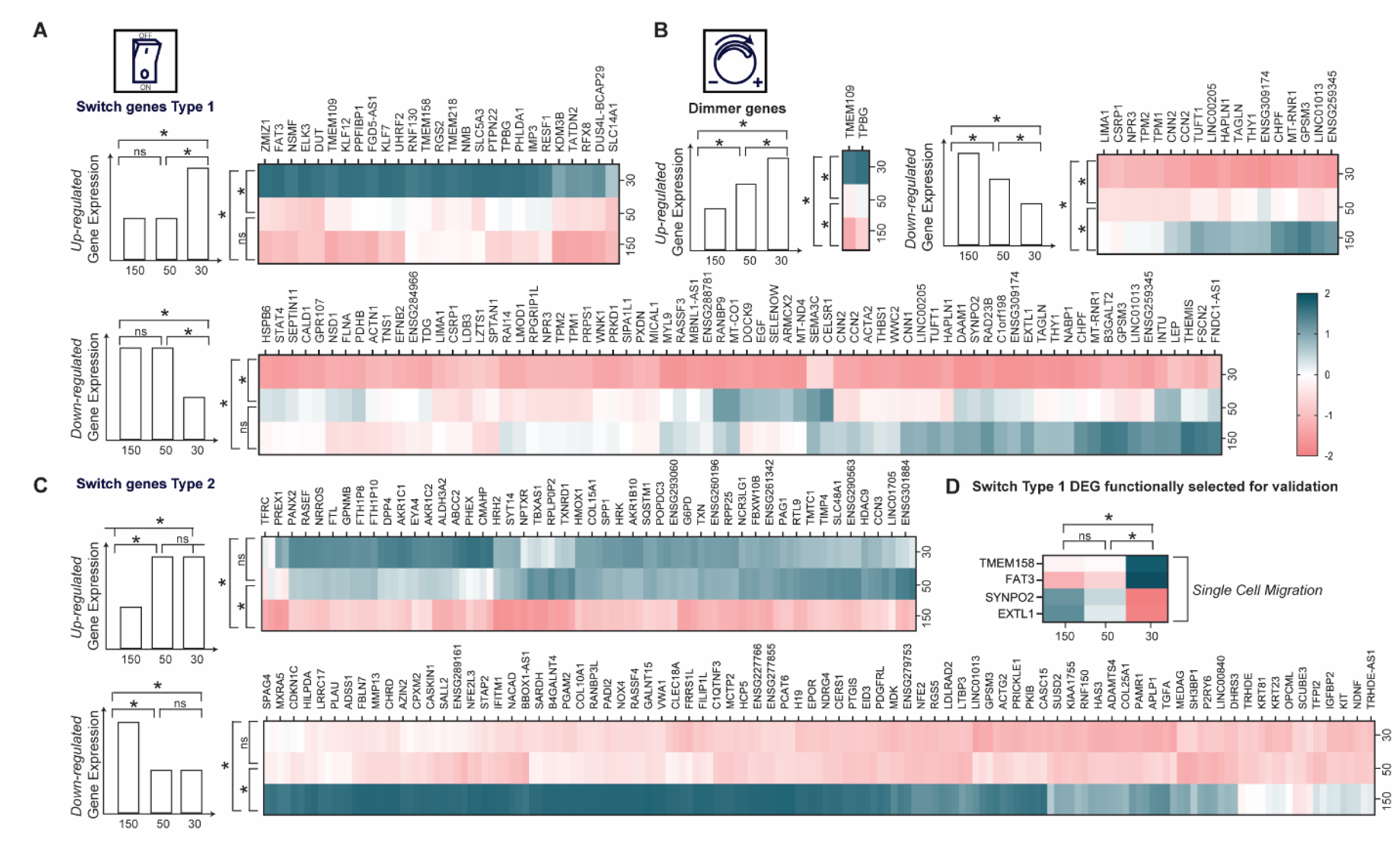
Identification of confinement-responsive Switch and Dimmer genes in GlaMS. (A) Type 1 Switch genes, defined as genes significantly up- or downregulated in 30 µm versus 150 µm and 30 µm versus 50 µm GlaMS, but not significantly altered between 50 µm and 150 µm GlaMS. (B) Dimmer genes, defined as genes significantly differentially expressed across all three pairwise comparisons. (C) Type 2 Switch genes, defined as genes significantly altered in both 30 µm versus 150 µm and 50 µm versus 150 µm GlaMS, but not significantly different between 30 µm and 50 µm GlaMS. (D) Selected Type 1 Switch genes, including TMEM158, FAT3, SYNPO2, and EXTL1, identified for subsequent validation based on their confinement-responsive expression patterns and potential relevance to confined single cell migration. All heat maps were made using transcript-per-million (TPM) values-based Z- scores. Statistical significance (*) was determined using adjusted p ≤ 0.05 and were further classified as strongly regulated (|log₂FC| ≥ 1) or mildly regulated (0 < |log₂FC| < 1), with positive and negative log₂FC values indicating upregulation and downregulation, respectively. ns is not significant.

### Microchannel assays confirm post-confined migration TMEM158 upregulation

Of all the Switch Type 1 DEGs, four candidate genes with single cell migration-related functions **(Fig. 6D)** were selected for further analysis with the goal of identifying previously uncharacterized regulators of confinement. TMEM158 (upregulated) has been reported to induce epithelial-to-mesenchymal transition (EMT) via activation of the PI3K/AKT pathway ^[51–54]^; FAT3 (upregulated), an atypical cadherin, regulates cell migration and polarity and localizes to the leading edge of migrating cells^[55–58]^; SYNPO2 (downregulated), an actin-binding protein, promotes migration via Arp2/3-dependent lamellipodia formation^[59,60]^; and EXTL1 (downregulated), a glycosyltransferase involved in heparan sulfate biosynthesis, plays a role in dendritic cell migration with knockout models demonstrating impaired migration^[61]^. Of these four, TMEM158 emerged as a particularly compelling candidate. Although it has not been studied in the context of confinement, it has been shown to regulate cancer cell migration in confining microenvironments, as its knockdown was found to reduce migration through short 8 µm transwell pores^[54]^.

To better understand confinement-dependent expression dynamics of these shortlisted candidates in an environment with lower throughput but more straightforward imaging, microchannel assays were deployed to compare unconfined cells (found in the entrance reservoir) to post-confinement cells (in the exit reservoir) **(Fig. 7A)**. Microchannel cross-sectional areas of 30 µm² and 100 µm² were specifically chosen to mimic the average pore areas found within 30 µm and 50 µm GlaMS (approximately 25 µm² and 100 µm², respectively) **(Fig. 1I).** Immunofluorescence revealed a confinement-sensitive upregulation of both cellular **(Fig. 7B, C)** and nuclear **(Fig. 7B, D)** TMEM158 expression, mirroring the upregulation of TMEM158 mRNA levels found in the GlaMS and validating the GlaMS as a tool for identifying confinement-sensitive genes.

**Figure 7:**
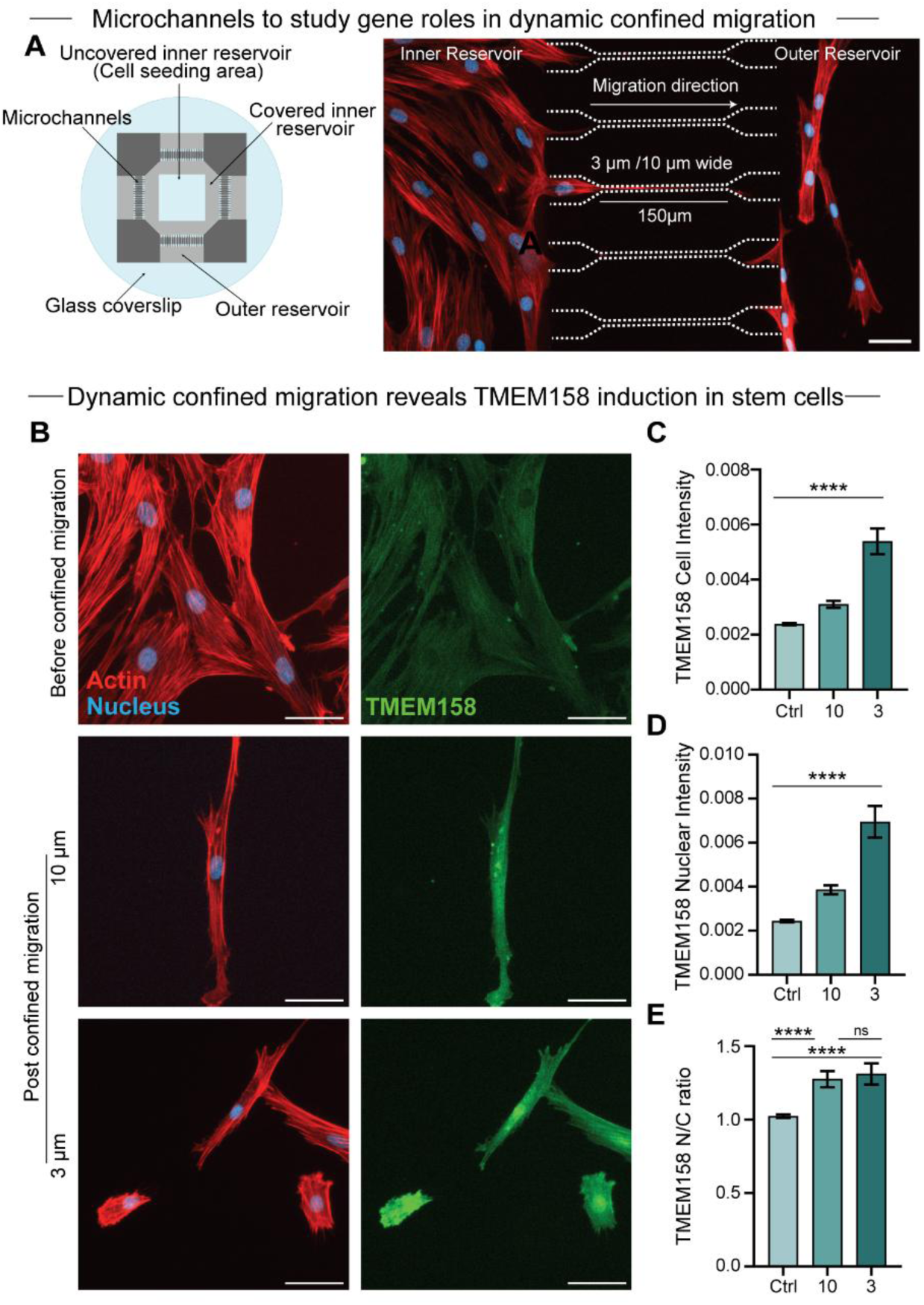
Confined migration increases expression of TMEM158 in stem cells. A) (Left) Microchannel chip schematic. (Right) Representative image demonstrating cells migrating through 150 µm long channels, 3 µm or 10 µm wide without any chemoattractant. B) Representative images of cells before and after confined migration. Scale bar = 50 µm. C) Quantification of TMEM158 mean intensity in cells post-confined migration. D) Quantification of nuclear mean intensity of TMEM158 in cells post-confined migration. E) Quantification of nuclear to cytoplasmic ratio of TMEM158 in cells post-confined migration. N=3, n=209, 90, 115, respectively. All graphs have error bars represented as 95% confidence interval. Kruskal-Wallis test with Dunn’s multiple comparison test for statistical significance. *P <= 0.05, **P <= 0.01, ***P <= 0.001, ****P <=0.0001.

Consistent with RNA-seq results, FAT3 expression also increased following confined migration in both 3 µm and 10 µm channels **(Fig. S12A-D**). Unexpectedly, both SYNPO2 and EXTL1, identified as downregulated *“Switch”* genes in the GlaMS, showed increased protein expression following confined migration (**Fig. S13, Fig. S14**). To understand this discrepancy, we performed qPCR using gene specific primers **(Fig. S13E, Fig. S14E)** using a TRAP (Trap-based Recovery After Permeation) microchannel system which allows extraction of low volume lysates from post-confined migration cells^[62]^. qPCR of these lysates confirmed the mRNA downregulation found in the RNA-Seq from GlaMS. Differences between mRNA and protein expression levels of certain genes are well documented in the literature, as protein output is further regulated by translation efficiency, protein stability, and degradation rates^[63]^. Temporal delays between transcriptional changes and protein turnover, as well as post-transcriptional and post-translational regulation, can further decouple RNA and protein readouts in cells^[64]^.

### Loss of TMEM158 hinders confined migration

To move beyond expression-based association and test whether TMEM158 plays a functional role in confinement adaptation, we performed loss-of-function studies using siRNA-mediated knockdown in hMSCs prior to microchannel migration assays **(Fig. 8A)**. TMEM158 knockdown was first confirmed by qPCR, which showed a nearly 90% reduction in mRNA levels relative to control **(Fig. 8B)**. Knockdown cells were then seeded into chips containing 3 µm and 10 µm microchannels for 48 hours. Significantly fewer cells were able to successfully migrate to the exit reservoir following TMEM158 knockdown, particularly under the more confining condition **(Fig. 8C,D)**. Importantly, TMEM158 knockdown did not significantly alter 2D migration speed, indicating that the reduced migration observed in microchannels was not due to a general motility defect **(Fig. 8E)** and suggesting that TMEM158 is specifically required for efficient adaptation to confined environments.

**Figure 8:**
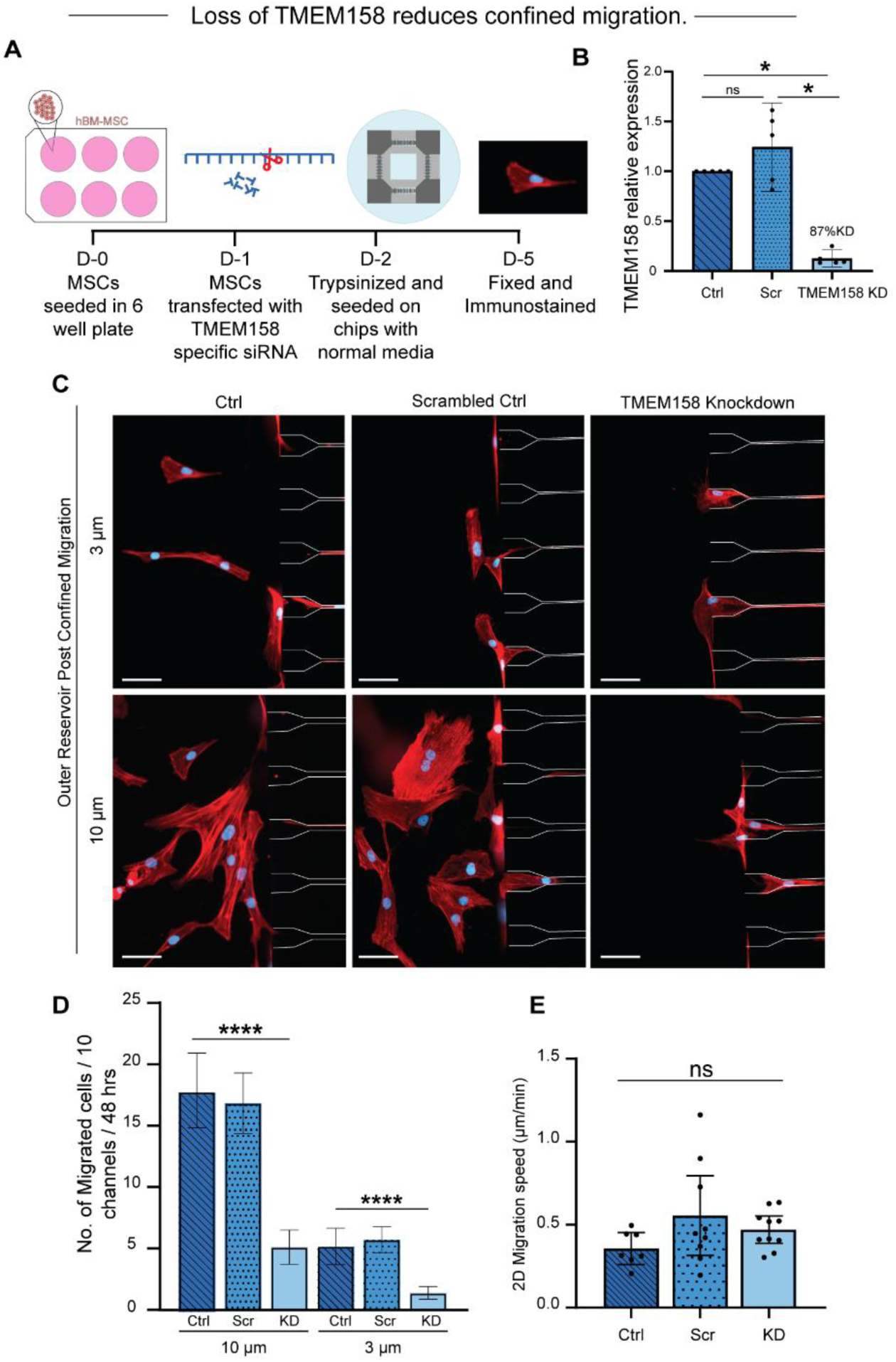
TMEM158 depletion compromises the ability of stem cells to migrate effectively under confinement. (A) Schematic of experimental workflow. hMSCs were seeded, transfected with TMEM158 specific siRNA or scrambled siRNA for 24 hours and subjected to confined migration assays in microchannels, followed by fixation and immunostaining. (B) qPCR validation of TMEM158 knockdown efficiency relative to control (Ctrl) and scrambled siRNA (Scr), showing ∼87% reduction in expression. (C) Representative immunofluorescence images of hMSCs post-migration through 3 µm and 10 µm wide microchannels for 48 hours. Cells were stained for F-actin (red) and nuclei (blue). Scale bars: 50 µm. (D) Quantification of the number of cells successfully migrating through 3 µm and 10 µm channels over 48 hours. N = 2, n = 300. (E) Quantification of 2D migration speed (µm/min). N = 2, n = 30. All graphs have error bars represented as 95% confidence interval. Kruskal-Wallis test with Dunn’s multiple comparison test done for statistical significance. *P <= 0.05, **P <= 0.01, ***P <= 0.001, ****P <=0.0001.

## Conclusion

This study demonstrates that non-annealed glass microparticle scaffolds (GlaMS) provide a scalable, physiologically relevant platform for long-term 3D confinement of cells. By mimicking *in vivo* confinement and allowing straightforward retrieval of substantially larger confined cell populations than conventional platforms, GlaMS overcomes the throughput and recovery limitations of existing models. LOVAMAP, a 3D pore segmentation tool, was used to predict the degree of confinement experienced by cells migrating through GlaMS. These dimensions were then compared with tissue interstitial spaces in *ex vivo* tissues to identify the sphere size that would best mimic *in vivo* confinement. Stem cells placed in these scaffolds show a strong migratory response, actively moving through both doors and pores and causing nuclear compression typical of confinement both *in vivo* and *in vitro*. Sustained confinement in GlaMS induces persistent morphological changes, accelerates migration, alters YAP signaling, and enhances osteogenic priming in hMSCs. Transcriptomic analyses further revealed confinement degree-dependent gene expression shifts and enabled the identification of confinement-responsive genes, offering new targets for mechanobiology research. The transmembrane protein TMEM158 was found to be heavily upregulated in response to physiological confinement, then further validated in microchannel assays. siRNA-induced TMEM158 knockdown resulted in a significant reduction in stem cell confined migration, suggesting that it plays a key role in both responding to confinement and equipping cells for confined migration. Together, these findings position GlaMS as a versatile tool for dissecting confinement-mediated mechanotransduction and for preconditioning therapeutic stem cells to improve regenerative outcomes in the future.

## Materials and methods

### Quantification of pore and door dimensions in GlaMS using LOVAMAP

Houdini by SideFX was used to generate simulated scaffolds of our three GlaMS groups: high confinement (30 µm diameter particles; N = 3 scaffolds), moderate confinement (50 µm diameter particles; N = 3 scaffolds), and unconfined (150 - 210 µm diameter particles; N = 10 scaffolds). Glass particles were simulated as rigid bodies under Newtonian dynamics, per Houdini. For 30 µm and 50 µm groups, monodisperse spheres **(Fig. 1J)** of the corresponding size were dropped into a 600 µm x 600 µm x 600 µm container as previously described^[23]^. For the 150-210 µm group, polydisperse spheres were randomly generated according to a distribution of experimental particle sizes **(Fig. 1J)** then dropped into a 1000 µm × 1000 µm × 1000 µm container. LOVAMAP software was run on each domain to identify doors and 3D pores, where doors represent the narrowest regions in space between neighboring 3D pores and 3D pores represent the widest pockets of space^[23]^. LOVAMAP output included void space descriptors, including door diameter (µm), 3D pore volume (picolitre), and diameter of the largest enclosed sphere (LES) in a 3D pore (µm) ^[23,38]^. Door diameter and LES were used to represent the narrow and wide dimensions through which cells could migrate within a scaffold’s void space. To compare these values with physiological confinement, width dimensions of tissue tracks such as those found in adipose networks, perineural spaces, and collagen-dense regions were compiled from published intravital imaging datasets^[1]^. The corresponding cross-sectional areas of these tissue tracks were calculated and overlaid with the LOVAMAP-derived pore and door size distributions, as shown in **Figure 1I**.

### Cell culture

Human bone marrow-derived mesenchymal stem cells (hBM-MSCs; CD105⁺, CD166⁺, CD44⁺, CD90⁺, CD73⁺, CD14⁻, CD34⁻, CD45⁻; Lonza, PT-2501) were expanded in T75 tissue culture flasks using Dulbecco’s Modified Eagle Medium high glucose (DMEM; Gibco, 10569) supplemented with 10% fetal bovine serum (FBS; Gibco, 26140079) and 1% penicillin-streptomycin (Pen/Strep; Gibco, 15140-122). RPE1-NLS-GFP cells were expanded in T25 tissue culture flasks using Dulbecco’s Modified Eagle Medium F12 (Gibco, 11320033) supplemented with 10% fetal bovine serum (FBS; Gibco, 26140079) and 1% penicillin-streptomycin (Pen/Strep; Gibco, 15140-122). Passage 12-14 of RPE1-NLS-GFP was used for all experiments. All cells were maintained at 37 °C in a humidified atmosphere of 5% CO₂ and passaged at 80-85% confluency. Subculturing was performed by detaching cells with 0.05% Trypsin–EDTA (Thermo Fisher Scientific, 15400054) and reseeding at a density of 5,000 cells cm⁻².

### Mouse tissue immunofluorescence

Liver and quadriceps muscles were harvested from 7-week-old white coat ICR mice following euthanasia. Prior to fixation, liver were incubated with heparin (30 min, 4 °C, with gentle shaking) to reduce residual red blood cells. Tissues were then fixed in 4% paraformaldehyde (PFA) overnight at 4 °C with shaking. Post-fixation, samples were washed three times in 1× PBS (1 hour each wash, 4 °C). Tissues were embedded in 4% low-melting-point (LMP) agarose in PBS. Livers were sectioned into 250 µm slices using a vibratome at a speed of 0.1 mm/s and stored in 1× PBS at 4 °C. Quadriceps muscles were sectioned either longitudinally or transversely. Tissue sections were permeabilized with 500 µL of Ce3D permeabilization buffer at 4 °C with rocking. For immunostaining, liver sections were incubated with primary antibodies: mouse monoclonal anti-Glucose Transporter GLUT1 (Santa Cruz, sc-377228, 1:100) and anti-Collagen I (Santa Cruz, sc-293182, 1:100). Following primary antibody incubation, sections were washed at room temperature with shaking using 1X PBS. Secondary staining was performed using anti-mouse Alexa Fluor 640 (1:500), phalloidin Alexa Fluor 568 (1:500), and hoechst 33342 (1:2000). After staining, tissue slices were mounted on glass slides and imaged using an Olympus FV3000 NIR confocal microscope.

### GlaMS assembly, cell seeding, harvest and imaging

Soda-lime glass microparticles of diameters 31-34 µm, 48-51 µm (Cospheric), and 150-210 µm (Polysciences) were sterilized, plasma-treated, and coated overnight with rat-tail type I collagen (100 µg/mL). For 3D confinement in GlaMS, hBM-MSCs were seeded onto collagen-coated microparticles at a density of 500 cells/µL of beads and allowed to attach overnight. Cell-bearing microparticles were subsequently transferred into 24-well Transwell inserts (8 µm pore size) and packed by gentle vertical tapping to form GlaMS. Cells were cultured for 3 or 7 days, with medium replaced every 2 days. Following confinement, cells were recovered by treatment with 0.25% trypsin-EDTA and replated onto collagen-coated glass coverslips for downstream immunofluorescence analysis. For RNA sequencing, cells were lysed directly within GlaMS using TRIzol after 7 days of confinement. Detailed procedures for microparticle preparation, cell seeding, GlaMS assembly, cell recovery, live imaging, and RNA isolation are provided in the SI Appendix, Extended Materials and Methods.

### Immunofluorescence

Cells on the glass coverslips were rinsed with 1X PBS and fixed in 3.7% paraformaldehyde for 20 min at room temperature (RT), followed by three washes in 1XPBS (5 min each). Cells were permeabilized with 0.1% Triton X-100 in PBST (PBS containing 0.1% Tween-20) for 10 min, then washed three times in PBST (3 min each). Non-specific binding was blocked using 1% bovine serum albumin in PBST for 30 min at RT. Primary antibodies: YAP1 (Rabbit polyclonal, 1:100, Invitrogen, PA1-46189) and RUNX2 (Mouse monoclonal, 1:100, Abcam, ab76956) and TMEM158 (Rb Polyclonal, 1:200, Invitrogen, PA5-84968) were diluted in 1% BSA/PBST and incubated overnight at 4 °C. After three PBST washes (5 min each), samples were incubated with secondary antibodies for 2 h at 37 °C: goat anti-mouse CF640 (1:1000, Biotium, 20175), or goat anti-rabbit CF488 (1:1000, Biotium, 20019). For F-actin visualization, F568 Phalloidin (1:100, Biotium, 00044) was included during secondary incubation. Nuclei were counterstained with Hoechst 33342 (1µg/mL, Invitrogen, H3570) for 10 min at RT, followed by three PBS washes. Coverslips were mounted using Fluoromount and dried overnight at 4 °C prior to imaging. All samples were imaged immediately after preparation.

### Bulk RNA sequencing and data analysis

Prior to sequencing, RNA integrity (RIN) was evaluated using high-sensitivity RNA electrophoresis (TapeStation). All RIN values were greater than 8. Standard bulk sequencing was performed on the NovaSeq X Plus sequencing platforms (NovogeneAIT Genomics Singapore Pte Ltd). RNA sequencing data was processed using the nf-core/differentialabundance pipeline, which includes quality control, annotations, and differential expression analysis using DESeq2. For the volcano plots, differentially expressed genes (DEGs) were identified using an adjusted *p* value ≤ 0.05 and a fold change of ≥ 2. For the identification of switch and dimmer genes, all significant genes with adjusted *p* value ≤ 0.05 were further classified according to the magnitude and direction of expression change as strongly upregulated (log₂FC ≥ 1), mildly upregulated (0 < log₂FC < 1), mildly downregulated (−1 < log₂FC < 0), or strongly downregulated (log₂FC ≤ −1). For heatmap visualization of gene expression across experimental conditions, transcript-per-million (TPM) values were averaged across five biological replicates for each condition. Mean TPM values were then used to calculate Z scores. Hierarchical clustering was performed using TBTools, in which genes in each row was hierarchically clustered. Volcano plots and enrichment bubble plots were generated using GraphPad Prism. Gene enrichment analysis was performed using WebGestalt 2024 with Gene Ontology (GO) and Kyoto Encyclopedia of Genes and Genomes (KEGG) gene sets for *Homo sapiens*.

### Image acquisition and morphological analysis

For fixed, fluorescently labeled samples, images were acquired on a Zeiss Celldiscoverer 7 widefield epifluorescence microscope using the 10X objective. Time-lapse imaging was performed on an Andor BC43 microscope in widefield fluorescence and transmitted light modes under environmental control (37 °C, 5% CO₂). Images were processed in Cell Profiler. Cells (primary object) and nuclei (secondary object) were segmented from actin (phalloidin) and Hoechst channels, respectively. Morphometric parameters included cellular area and aspect ratio (actin channel), and nuclear area and aspect ratio (Hoechst channel) were subsequently calculated. Mean fluorescence intensities of transcription factors (e.g., YAP and RUNX2) were extracted for cell, nucleus, and cytoplasm (tertiary object). The N/C ratio was calculated as

N / C ratio = Mean_n_ / Mean_c_

Where Mean_n_ is mean intensity of nucleus and Mean_c_ is mean intensity of cytoplasm.

### Characterization of cell migration of replated cells

hMSCs were confined in GlaMS for 3 days and subsequently replated on collagen-coated glass coverslips for 1 hour. Live-cell imaging of migration was performed using the BC42 benchtop widefield microscope with a 10X objective. Images were acquired every 10 minutes for a total duration of 6 hours following the initial 1-hour attachment period. Cell migration was quantified using the Manual Tracking plugin in FIJI, and the resulting XY coordinates were analyzed with the Ibidi Chemotaxis and Migration Tool to determine migration velocity and generate trajectory rose plots.

### siRNA-mediated knockdown and qPCR validation

hMSCs were transfected with TMEM158-targeting siRNA or a non-targeting scrambled control (Silencer™ Negative Control No. 4 siRNA, Thermo Fisher Scientific) using a lipid-based transfection reagent (Lipofectamine RNAiMAX, Thermo Fisher Scientific) at a final concentration of 50 nM, according to the manufacturer’s protocol. Transfection was performed for 24 hours, after which media was replaced with fresh growth media. TMEM158 targeting siRNA sequence: (Sense) CUGAAGUCUAGGCAGCAGAAUGCTT. Total RNA was extracted and reverse transcribed into cDNA using iScript cDNA Synthesis Kit (Bio-Rad). Quantitative PCR was performed using SSO Advanced SYBR Green Supermix (Bio-Rad) with the following primers: Forward: CCCGGGCCGCTGTGGCTGCAG; Reverse: GGCTGCGGTGGTGCTGGCGGTG.

GAPDH was used as the housekeeping control to normalize gene expression. Knockdown efficiency was validated by qPCR. Following transfection, cells were trypsinized and replated into 3 µm and 10 µm-wide microchannel devices, where they were allowed to migrate for 48 hours before fixation and staining for F-actin and DAPI.

### Statistical analysis

All graphs were reported as mean with 95% confidence interval and analyzed in GraphPad Prism (10.0). Normality was determined using the Shapiro-Wilk test. For comparisons between more than 2 groups, Kruskal-Wallis test with Dunn’s multiple comparisons (for non-normally distributed data) was used. Statistical significance between two groups was assessed using an unpaired t-test with Welch’s correction. P values less than 0.05 were considered statistically significant. ns is P > 0.05, * is P <= 0.05, ** is P <= 0.01, *** is P <= 0.001, **** is P <=0.0001.

## Acknowledgements

The authors acknowledge the support of the National Research Foundation of Singapore under an NRF Fellowship to Andrew W. Holle (NRFF13-2021-0114) and the support of the National Research Foundation, Singapore, under its Mid-sized Grant (NRF-MSG-2023-0001). The authors thank Jennifer Marlena for her immense help in creating the graphical abstract artwork.

## Data Availability

All raw RNA sequencing data have been deposited in the NCBI Sequence Read Archive (SRA) under BioProject accession number PRJNA1465086. Requests for further information and resources should be directed to the lead contact, Andrew W. Holle.

