## Supplementary Information and Extended Methods for "Rigid granular scaffolds reveal transcriptional responses to cellular confinement"

**This document includes:**

Extended materials and methods

Figures S1- S15

Legends for Supplementary videos 1-3

**Other supporting materials for this manuscript include the following:**

Videos 1 to 3

**Extended Materials and Methods**

***Cell culture:*** Commercially purchased Lonza PT-2501 contained hBM-MSCs isolated from bone marrow of the posterior iliac crests of normal volunteers between the ages of 18 and 45 years old. N denotes the number of independently performed experiments conducted using separately prepared scaffolds and cells from different passage numbers, whereas n denotes the total number of individual cells analyzed across these experiments. Cells were used between passages 4 and 8, which are considered early passages for cultured hBM-MSCs. MSC surface-marker expression and trilineage differentiation capacity within this passage range have previously been characterized by the manufacturer and in independent studies.

***Glass bead preparation and collagen coating* :** Precision spherical, monodisperse soda lime glass microparticles with diameters of 31-34 µm and 48-51 µm (Cospheric; S-SLGMS-2.5) and 150-210 µm (Polysciences; 05483-250) were used as granular scaffolds. Microparticles were composed of solid soda-lime glass (soda-lime silicate glass), which typically consists of ~75% silica (SiO₂), 14% sodium oxide (Na₂O), and 10% calcium oxide (CaO). Key material properties include a true particle density of ~2.5 g/cc, refractive index of ~1.5, annealing temperature of ~500 °C, and softening temperature of ~650 °C. Glass generally has a Young’s modulus of approximately 70 GPa. Particle size distribution was validated using brightfield images (**Fig. 1J**) to ensure accurate characterization of the size of the microparticles. Prior to cell culture, particles were sterilized by washing with 70% ethanol followed by three washes with sterile water. After drying overnight in an oven, 0.25 gm of particles were plasma-treated and coated with rat tail type I collagen (100 µg mL⁻¹; Gibco, Carlsbad, CA) and incubated overnight at 37 °C. The following day, collagen-coated particles were rinsed in phosphate-buffered saline (PBS), sterilized under UV light for 15 min prior to use. Beads were allowed to settle under gravity in 15 mL falcons for 1 min before resuspending in sterile cell culture media at 2 g/mL concentration. Similar to the glass microparticles, glass coverslips measuring 12 mm in diameter were cleaned with 70% ethanol, dried, and plasma-treated. The coverslips were subsequently coated with rat-tail type I collagen at 100 µg mL⁻¹ and incubated overnight at 37°C. The following day, the collagen-coated coverslips were rinsed with 1X PBS and sterilized under UV light for 15 min before use.

***Cell attachment and assembly of GlaMS*** ***(Fig. S7):*** For initial cell attachment, 100 µL of collagen-coated glass microparticles were seeded as a monolayer in Corning® 6-well ultra-low attachment plates (Corning, 3471). 1 mL of hMSCs at a concentration of 0.05 million cells/mL was added to the microparticle monolayer (500 cells/µL of beads) and incubated overnight to promote cell attachment. Attachment of the cells to the microparticles was verified microscopically after 24 hours **(Fig. S5C)**. The cell-bearing microparticles were transferred into 24-well Transwell® inserts (Corning, 3428) of pore size 8 µm using a 1 mL pipette tip. The bead and cell suspension was allowed to settle under gravity for 1 min, while gentle vertical tapping of the plate was used to facilitate random close packing of the microparticles. Subsequently, 1 mL of excess cell culture medium was added per well and replaced every 2 days to ensure adequate nutrient supply during confinement. The cells were maintained within the GlaMS at 37 °C in a humidified atmosphere containing 5% CO₂ for either 3 or 7 days.

***Cell harvest from GlaMS post-confinement*** ***(Fig. S8)****:* Following the confinement period, the cell-bearing microparticles were transferred from the transwell inserts into a 15 mL Falcon tube using a 1 mL pipette tip. The microparticles were allowed to settle under gravity for 1 min, after which the supernatant was carefully removed and transferred to a 24-well plate. The 24-well plate was incubated for 24 hours and examined for the presence of any live cells that may have detached during transfer under a phase contrast microscope. The cell-bearing microparticles were then washed with sterile 1X PBS. The tube was gently tapped to mix the suspension and remove any residual culture medium. The microparticles were allowed to settle for 1 min, after which the PBS supernatant was removed and transferred to a 24-well plate. This plate was also incubated for 24 hours and examined for the presence of any live cells that may have detached during washing under a phase contrast microscope. To detach the cells from the microparticles, 0.25% Trypsin-EDTA (Thermo Fisher Scientific, 15400054) was added, and the tube was gently tapped to ensure thorough mixing. The suspension was incubated at 37 °C for 5 min, after which 1 mL of cell culture medium was added to neutralize the excess trypsin. The microparticles were allowed to settle for 1 min, and the supernatant containing the detached cells was transferred to a fresh tube. This cell-recovery step was repeated twice with cell culture media to minimize cell loss. The pooled cell suspension was centrifuged at 1000 rpm for 3 min, and the resulting cell pellet was collected for replating. Following cell recovery, the microparticles were examined under phase contrast microscope to confirm that all cells had detached. The cell pellet harvested from the GlaMS was resuspended in 1 mL media and seeded onto the collagen-coated glass coverslips. Cells were allowed to attach and spread at 37°C for 3, 6, 12, or 24 hours before proceeding with immunostaining.

***Cell seeding and in situ harvest from GlaMS for RNA sequencing*** ***(Fig. 11A)****:* Cells were attached to glass microparticles and confined in GlaMS as previously described. For the RNA-Seq, hMSCs at a concentration of 0.1 million cells/mL was added to 200 µL of collagen-coated glass microparticles keeping the cell density constant at 500 cells/ µL of beads. After 7 days confinement in GlaMS excess culture medium was gently removed from the 24-well Transwell® inserts, and 1 mL of cold TRIzol™ reagent (Thermo Fisher Scientific, 15596026) was added directly to each GlaMS. The cell-bearing microparticles suspended in TRIzol™ were then transferred into a 1.5 mL microcentrifuge tube using a 1 mL pipette tip. The tube was gently tapped to thoroughly mix the components, facilitate complete lysis of the cells within the GlaMS, and was subsequently incubated on ice for 5 min. After the microparticles had settled during the incubation period, the supernatant containing the cell lysate, approximately 950 µL, was transferred to a fresh tube. The remaining microparticles were centrifuged at 1000 rpm for 3 min to densely pack the beads and recover any residual lysate. The remaining supernatant was collected and added to the tube containing the initial cell lysate. The microparticles were subsequently examined under a phase-contrast microscope to confirm that no cells remained attached. The collected cell lysates were stored at −80°C until RNA extraction. Each experimental replicate (N) was performed on a different day using cells from a different passage number between P4 and P8. RNA extraction from all five experimental replicates was performed on the same day. RNA was purified using silica-column purification with the Qiagen RNeasy Mini Kit (Qiagen, 74104), according to the manufacturer’s protocol for mammalian cells, and the purified RNA was stored at −80 °C.

***Cell seeding on top of GlaMS*** ***(Fig 2A-C)*** *:* For cell seeding on top of GlaMS, collagen-coated glass microparticles were first added to each well of a 96-well plate to a height of 1mm and gently tapped vertically to facilitate random close packing. 100 µL of hBM-MSCs at a concentration of 0.15 million cells/mL was then seeded onto the top surface of the packed GlaMS. Subsequently, 200 µL of cell culture medium was added to each well. Before imaging, the culture medium was supplemented with 1 µg/mL Hoechst 33342 (Thermo Fisher Scientific, 33342), and the cells were incubated for 15 min. The bottom of each well was then imaged to assess the presence of cells that had migrated through the GlaMS. Following imaging, the Hoechst-containing medium was gently removed and replaced with regular cell culture medium. This staining and imaging procedure was repeated on each imaging day.

***Live imaging of cell migration within GlaMS*** ***(Fig. 2D and Fig. S6A)****:* hBM-MSCs and RPE1-NLS-GFP cells were added at a concentration of 0.05 million cells/mL to 100 µL of collagen-coated glass microparticles in low-attachment plates as described before. For nuclear visualization of hBM-MSCs, siR-DNA (SpiroCHROME), a far-red nuclear dye, was added to the hBM-MSC culture medium at a final concentration of 50 nM. The following day, the siR-DNA-containing medium was gently removed and replaced with regular cell culture medium. PDMS wells measuring 1 mm in diameter and 2 mm in height were plasma-treated and attached to 15 mm glass coverslips. The microparticles bearing the attached, nuclear-stained cells or the RPE1-NLS-GFP cells were then transferred into the PDMS wells using a 1 mL pipette tip. The holders were gently tapped vertically to facilitate packing of the microparticles before live imaging. PDMS holders were placed in wells with excess media throughout the culture period. Live imaging was performed using an Olympus FV3000 confocal microscope equipped with a 30X silicone immersion objective (NA ~1.05). Silicone immersion oil (RI ~1.40) was used to reduce the refractive index gap. Imaging was performed at multiple z-stacks spanning 2 to 3 bead layers (~70 μm).

***Cell seeding and harvest for 2.5D control (Fig. S9A)****:* 100 µL of collagen-coated glass microparticles were seeded as a monolayer in Corning® 6-well ultra-low attachment plates (Corning, 3471). 1 mL of hMSCs at a concentration of 0.05 million cells/mL was added to the microparticle monolayer and incubated overnight to promote cell attachment. For the 2.5D curvature control experiment, the cell bearing microparticles were transferred to a 15 mL falcon tube and cells were harvested and replated using the same methodology as described previously.

*Microchannel confined migration:* Silicon wafers consisting of four arrays of microchannels with uniform dimensions of 10 µm in height, 150 µm in length, and widths of 3 and 10 µm were produced through a two-step photolithography process by the microfabrication core at the Mechanobiology Institute, NUS. Polydimethylsiloxane (PDMS) (SYLGARD 184, Dow Corning, Wiesbaden, Germany) was mixed at a 10:1 base-to-curing agent ratio and poured onto the microchannel silicon wafer. The mixture and wafer were vacuum degassed for 10 minutes to remove any trapped bubbles and placed in an oven to cure at 80 °C for 90 minutes. The PDMS was cooled for 30 minutes and then peeled from the wafer and cut into square chips with a central inner reservoir and outer reservoirs. The cut chips, along with 30 mm glass coverslips, were then treated with oxygen plasma (0.7 mbar, 300 W) for 1 minute before being attached and placed in an oven at 80 °C for 10 minutes to strengthen the surface bonding. The chips were placed in a six-well plate, and 100 µg mL−1 Rat Tail Type I Collagen (Gibco, Carlsbad, CA) was added carefully to the corners of the inner reservoir and the outer reservoirs and allowed to diffuse through the microchannels. The chips were vacuum degassed for 5 minutes and incubated at 37 °C overnight. Following overnight incubation, the chips were thoroughly rinsed with 1X PBS three times to remove excess collagen before being sealed with parafilm and stored at 4°C until needed. On the day of cell seeding, the microchannel chips were removed from 4°C, rinsed thoroughly with PBS once, and sterilized under UV light for 30 minutes. After UV sterilization, the PBS in the chips was aspirated completely and replaced with cell culture media. The chips were then transferred to the incubator for a minimum of 30 minutes to normalize the temperature to 37°C. 10 µl of hMSCs at a concentration of 1 million cells/mL was meticulously added to the corners of the inner reservoirs. After 4 hours, once the seeded cells were fully adherent, the chips were submerged in culture media to avoid any differences in hydrostatic pressure between the inner and outer reservoirs and to facilitate diffusion of media across microchannels. The culture media was changed every two days. After 4 days of confined migration, hMSCs were fixed and stained as previously described.

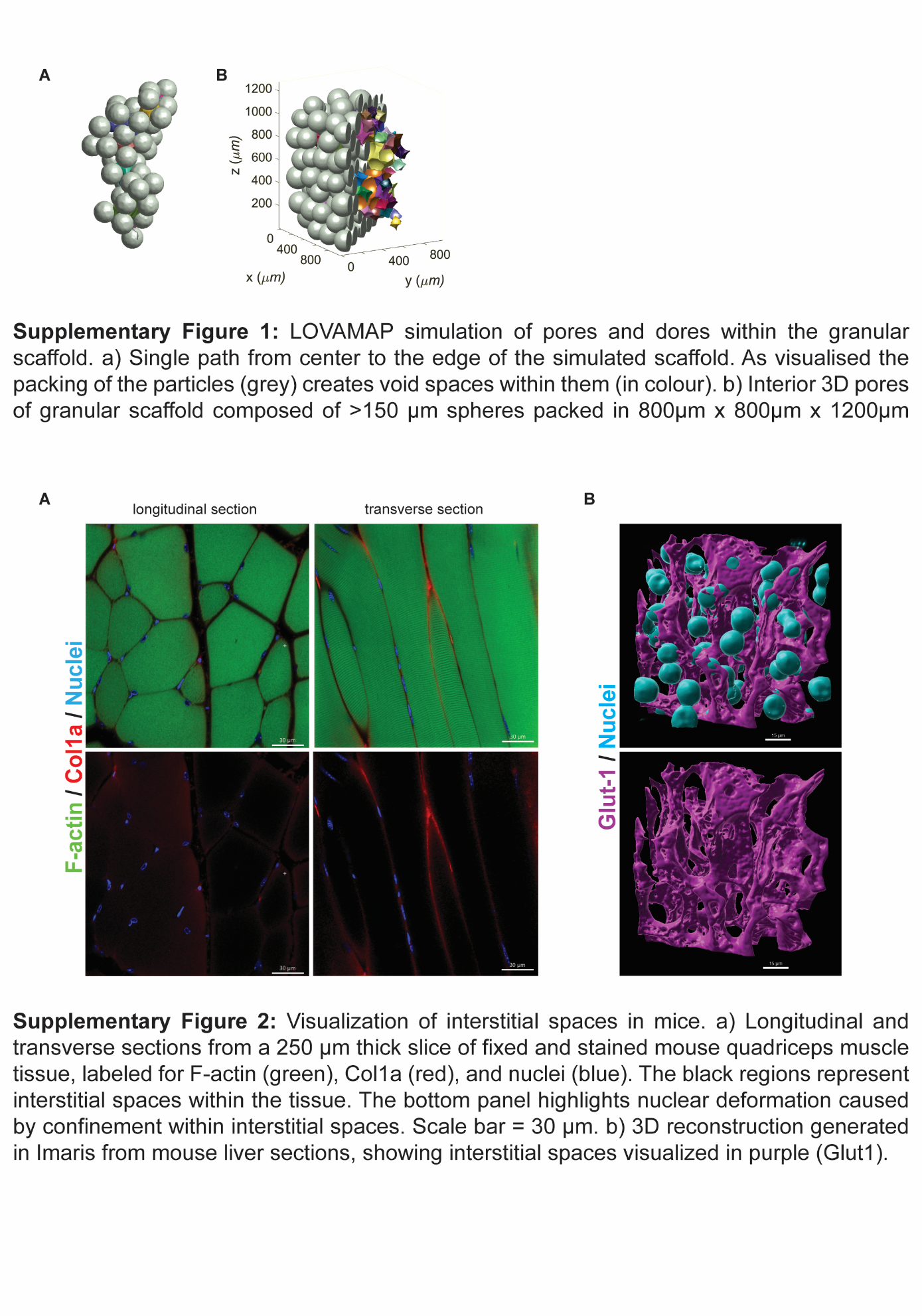

***Supplementary Figure 1: LOVAMAP simulation of pores and doors within the granular scaffold.*** *A) Single path from center to the edge of the simulated scaffold. As visualized, packing of the particles (gray) creates void spaces within them (in color). B) Interior 3D pores of granular scaffold composed of 150 μm spheres packed in 800 μm x 800 μm x 1200 μm holder.*

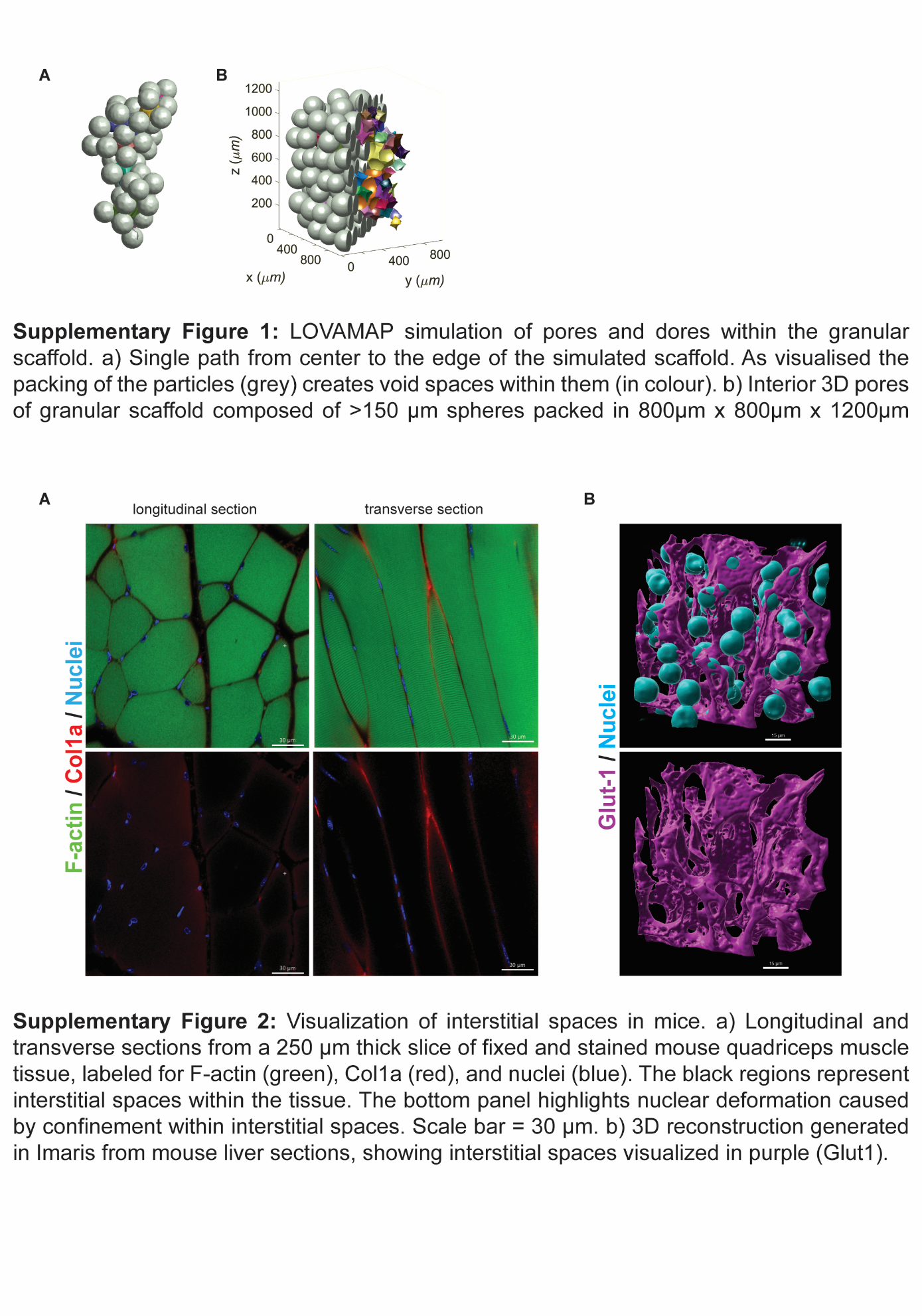

***Supplementary Figure 2: Visualization of interstitial spaces in mice.*** *A) Longitudinal and transverse sections from a 250 µm thick slice of fixed and stained mouse quadriceps muscle tissue, labeled for F-actin (green), Col1a (red), and nuclei (blue). The black regions represent interstitial spaces within the tissue. The bottom panel highlights nuclear deformation caused by confinement* *within interstitial spaces. Scale bar = 30 µm. B) 3D reconstruction generated in IMARIS from mouse liver sections, showing interstitial spaces visualized in purple (Glut1). Scale bar = 20 µm.*

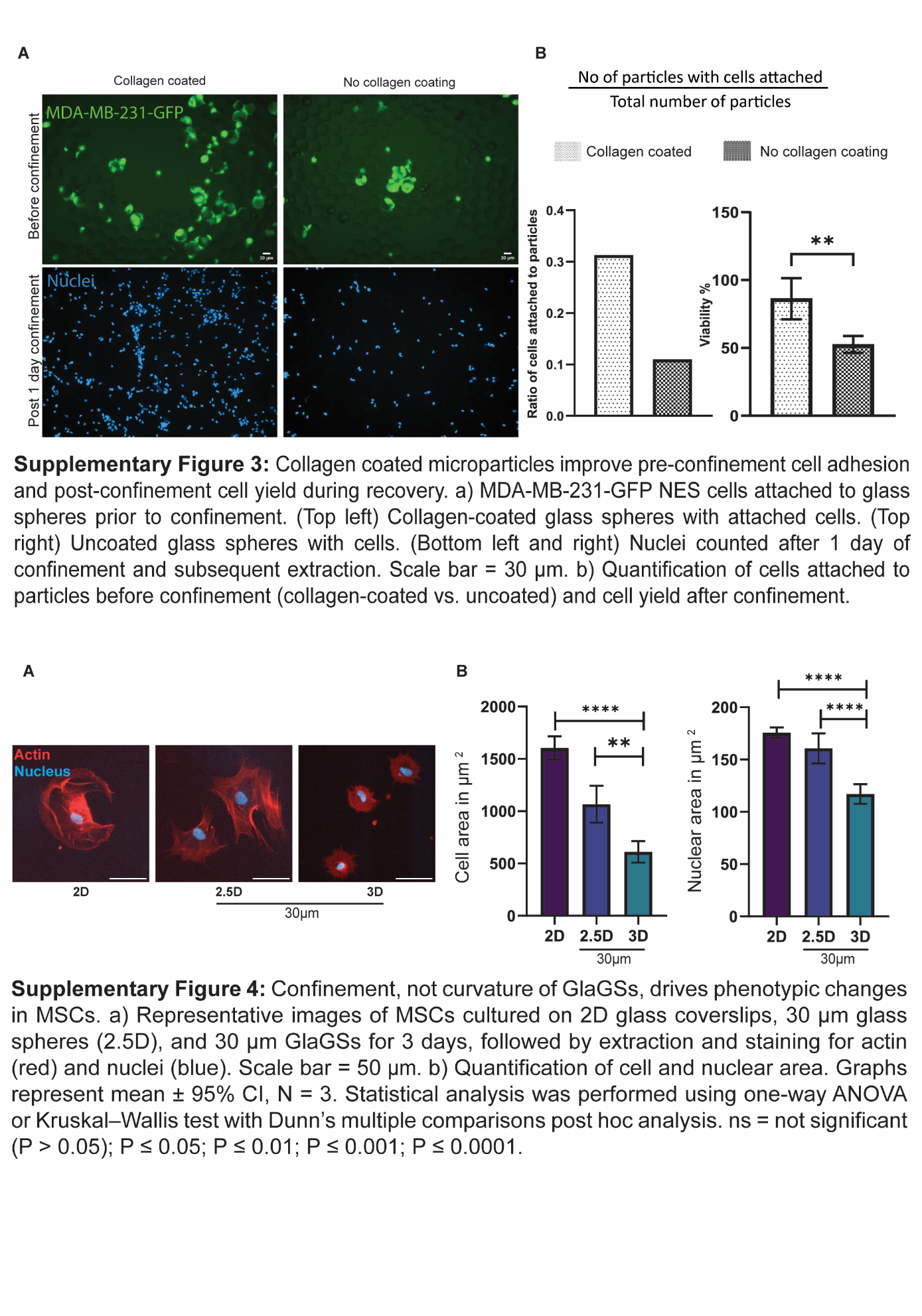

***Supplementary Figure 3: Collagen-coated microparticles improve pre-confinement cell adhesion and post-confinement cell yield during recovery.*** *A) MDA-MB-231-GFP NES cells attached to glass spheres prior to confinement. (Top left) Collagen-coated glass spheres with attached cells. (Top right) Uncoated glass spheres with cells. (Bottom left and right) Nuclei counted after 1 day of confinement and subsequent extraction. Scale bar = 30 µm. B) Quantification of cells attached to particles before confinement (collagen-coated vs. uncoated) and cell yield after confinement. Graphs have error bars represented as 95% confidence interval. Kruskal-Wallis test with Dunn’s multiple comparison test done for statistical significance. *P <= 0.05, **P <= 0.01, ***P <= 0.001, ****P <=0.0001.*

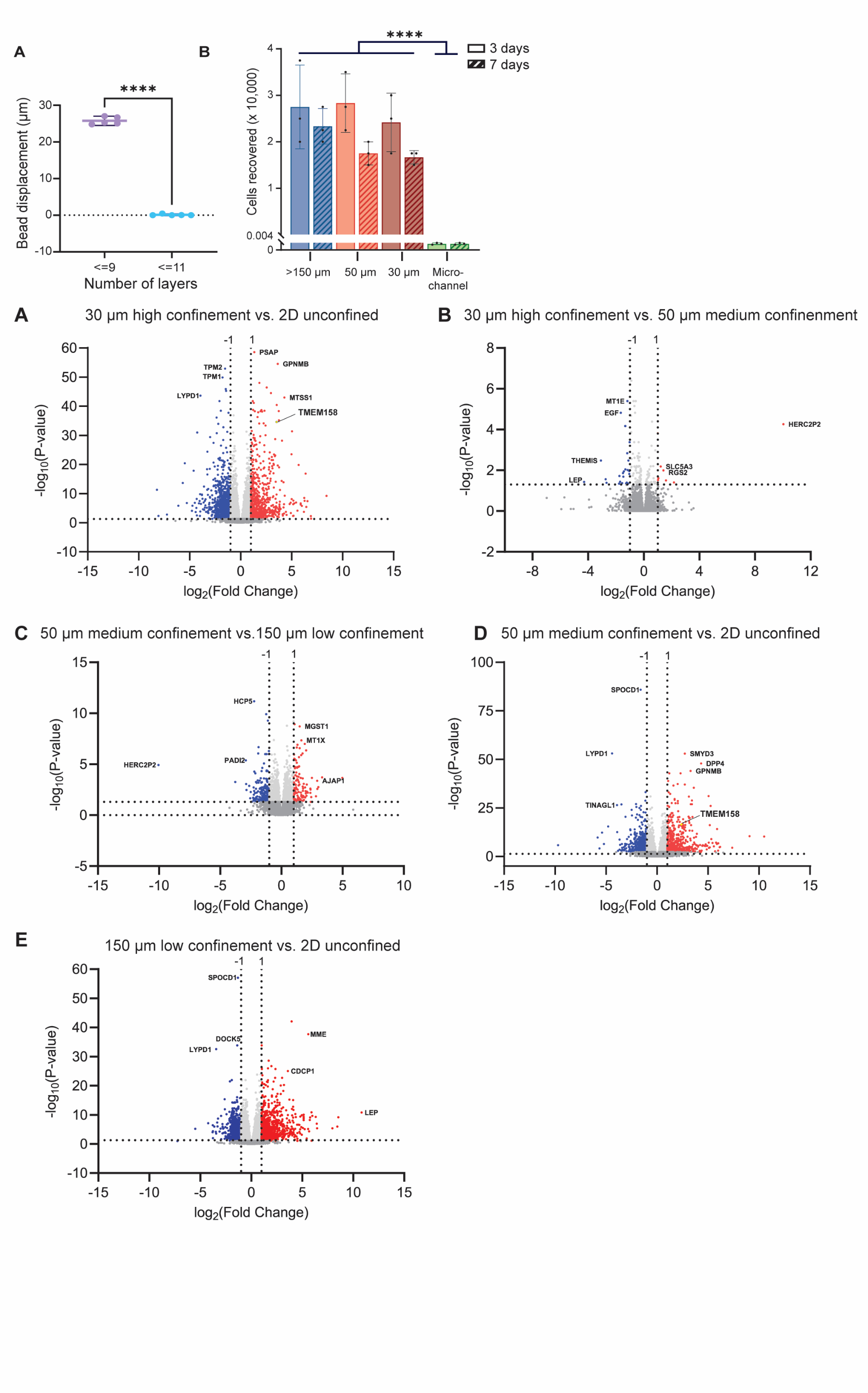

***Supplementary Figure 4: GlaMS enable long-term cellular confinement of large cell populations and facilitate straightforward recovery.*** *A) Confinement in GlaMS requires jammed packing of glass beads. Bead displacement graph at different bead layers. N = 2, n = 5. Error bars represent 95% confidence intervals. Unpaired t test with Welch’s correction was applied to calculate significance. B) Post-extraction, total cell counts demonstrate the high-throughput capacity of GlaMS compared to conventional in vitro confinement platforms such as microchannels. N = 3, Error bars represent 95% confidence intervals. Kruskal-Wallis test with Dunn’s multiple comparison test done for statistical significance. *P <= 0.05, **P <= 0.01, ***P <= 0.001, ****P <=0.0001.*

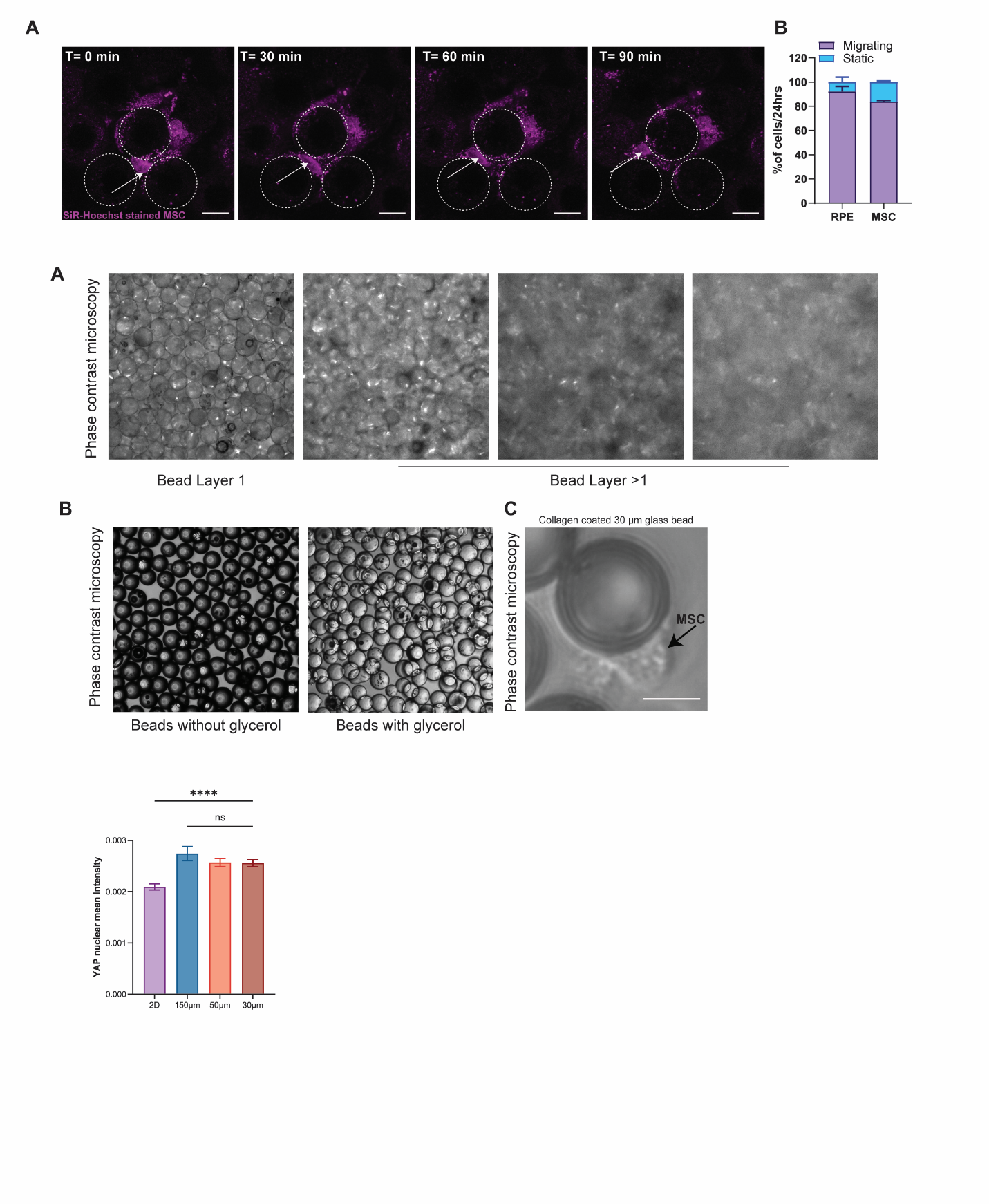

***Supplementary Figure 5:*** *A) Refractive index mismatch prevents imaging GlaMS beyond the first microparticle layer using conventional phase contrast microscopy. B) Refractive index matching using glycerol (RI =1.47) C) Phase contrast image of a P6 hMSC (black arrow) attached to a 30 µm collagen-coated glass bead in low attachment well plate, before packing to form GlaMS. Scale bar: 15 µm.*

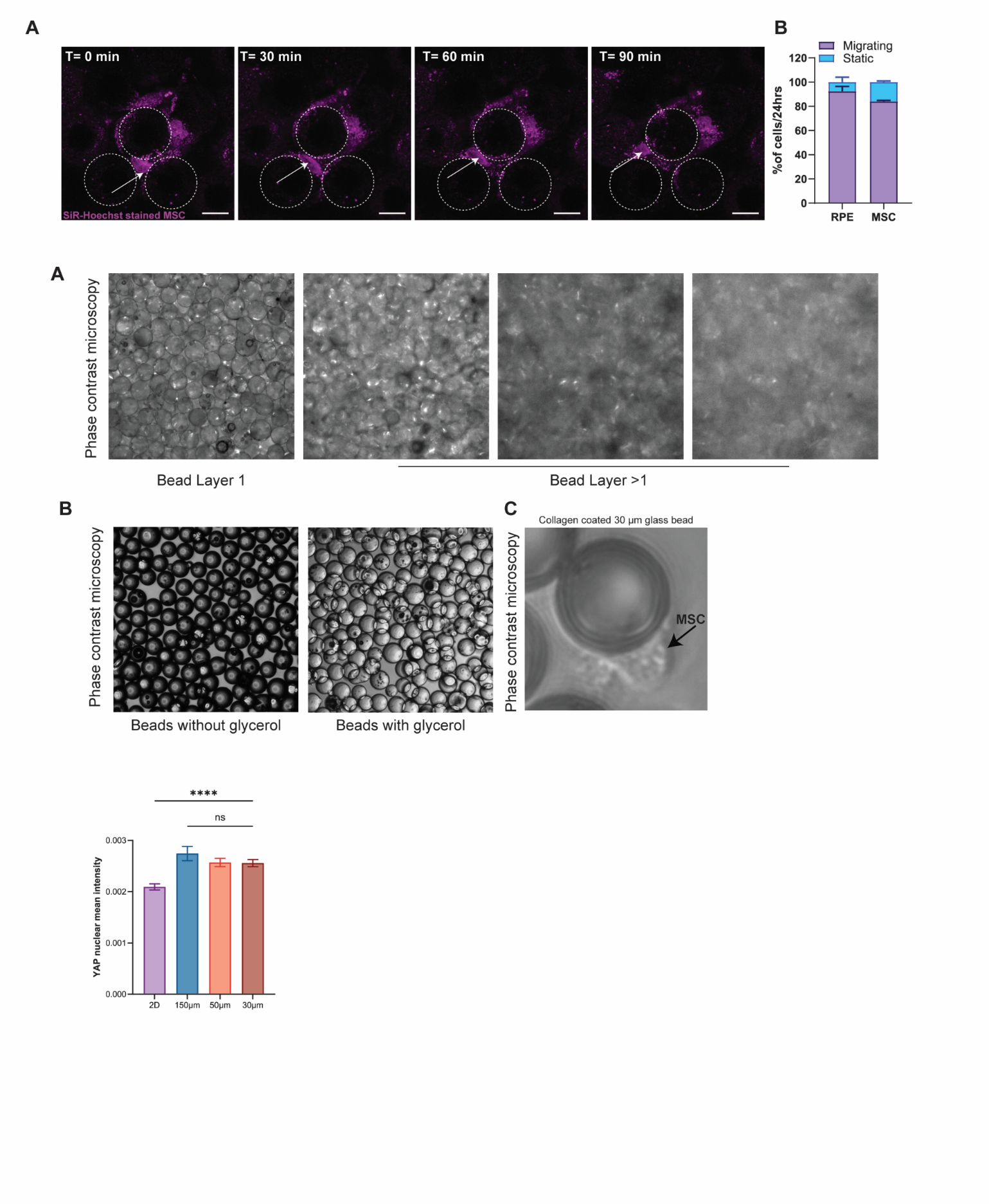

***Supplementary Figure 6: hMSCs undergoing active confined migration through interstitial spaces in GlaMS.*** *A) Time-lapse confocal images of SiR-DNA stained* *nuclei (magenta) in hMSCs migrating through interstitial voids formed by closely packed 30 µm GlaMS. Scale bars: 20 µm.* *B) Bar graph showing the percentage of migrating and static cells after 24 hours in GlaMS for RPE and hMSCs. Data are presented as mean with SD. N = 2, n = 40, 30 respectively.*

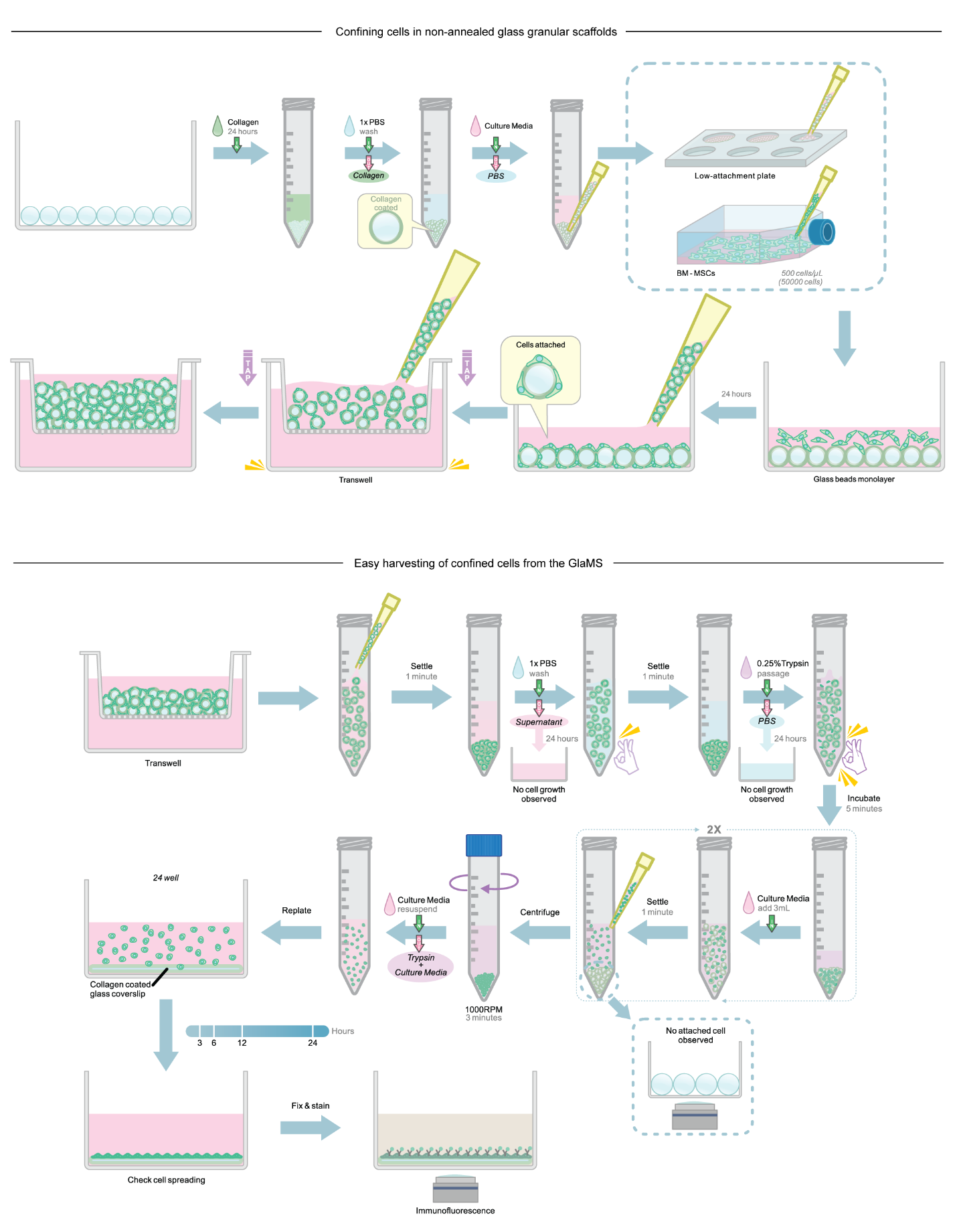

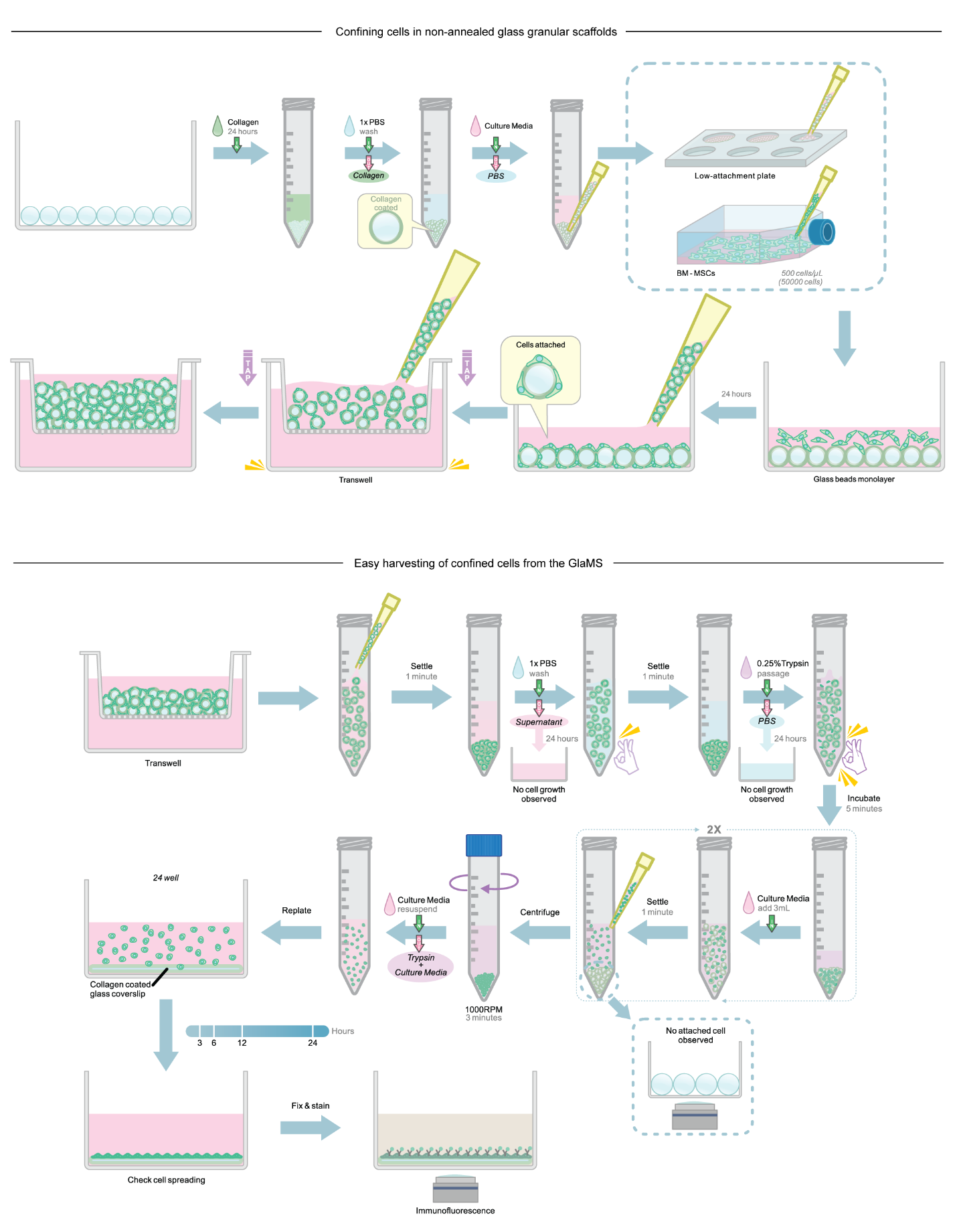
***Supplementary Figure 7:*** Methodology used to confine cells in non-annealed GlaMS.

***Supplementary Figure 8:*** Methodology used for easy retrieval of confined cells from GlaMS post-confinement

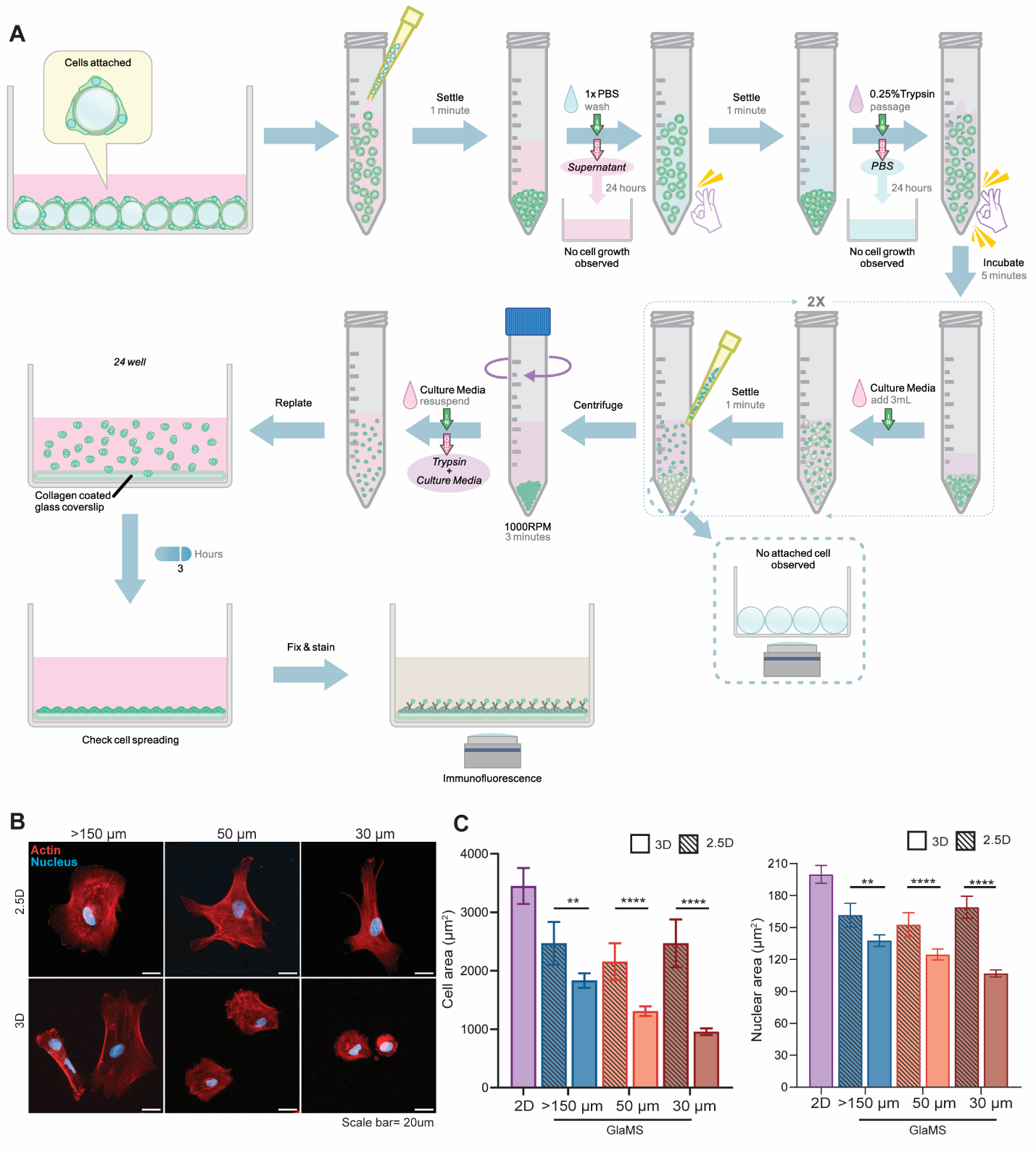

***Supplementary Figure 9: Confinement, not curvature, drives hMSC phenotypic changes in GlaMS.*** *A) Methodology used to culture and harvest cells from 2.5D bead surfaces. B) Representative images of hMSCs cultured on 150 µm, 50 µm and 30 µm glass spheres (2.5D), and 150 µm, 50 µm and 30 µm GlaMS for 3 days, followed by replating and staining for actin (red) and nuclei (blue). Scale bar = 20 µm. C) Quantification of cell and nuclear area. Graphs represent mean ± 95% confidence interval. Statistical analysis was performed using Kruskal-Wallis test with Dunn’s multiple comparisons post hoc analysis. *P <= 0.05, **P <= 0.01, ***P <= 0.001, ****P <=0.0001.*

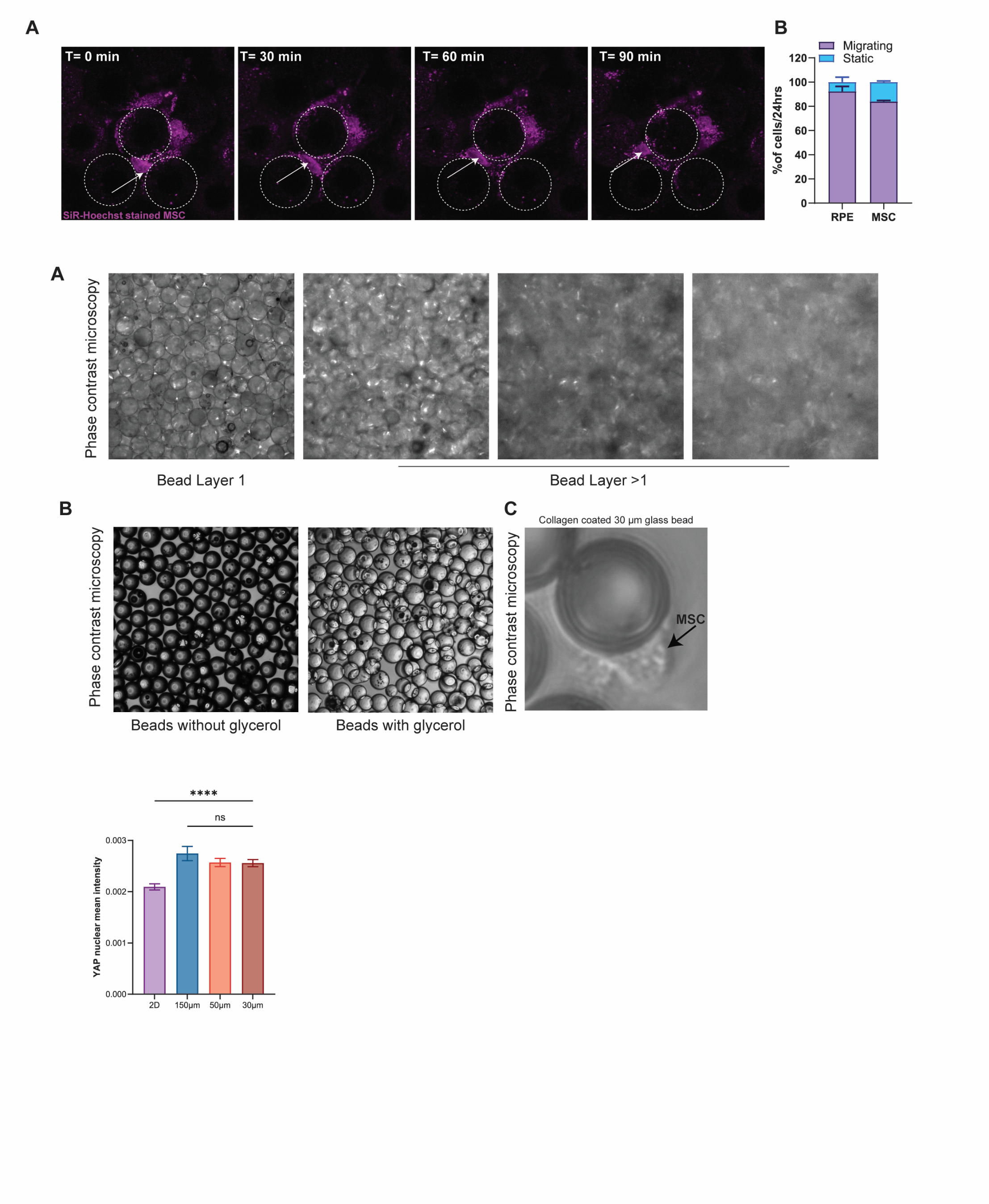

***Supplementary Figure 10: Quantification of YAP nuclear intensity across confinement conditions.*** *Bar graph showing YAP nuclear mean intensity in cells cultured in 2D and within GlaMS composed of 150 µm, 50 µm, and 30 µm beads. Data are presented as mean ± 95% confidence interval. N = 3, n = 197, 307, 338, 339, respectively. Statistical analysis was performed using Kruskal-Wallis test with Dunn’s multiple comparisons post hoc analysis. *P <= 0.05, **P <= 0.01, ***P <= 0.001, ****P <=0.0001.*

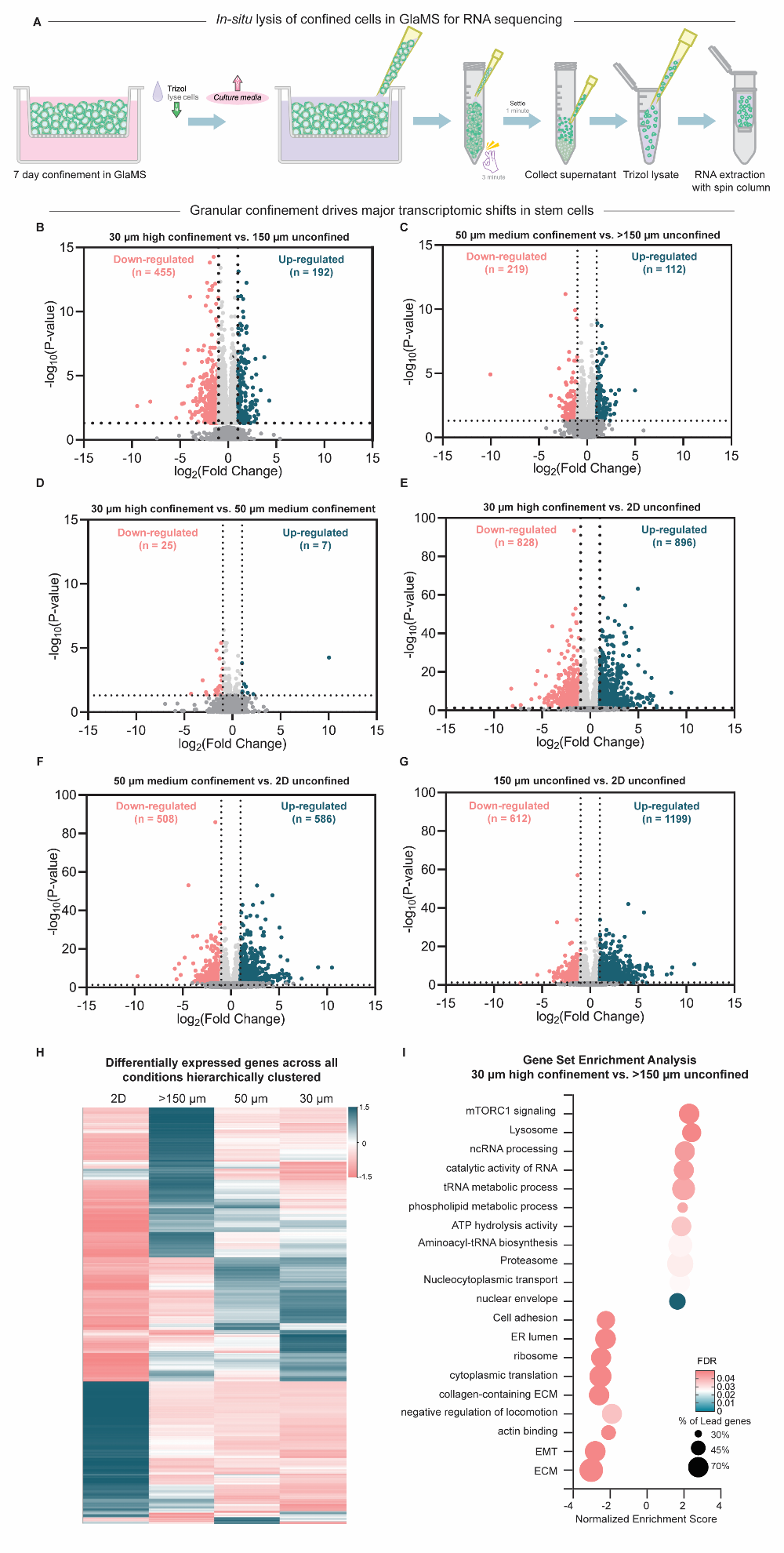

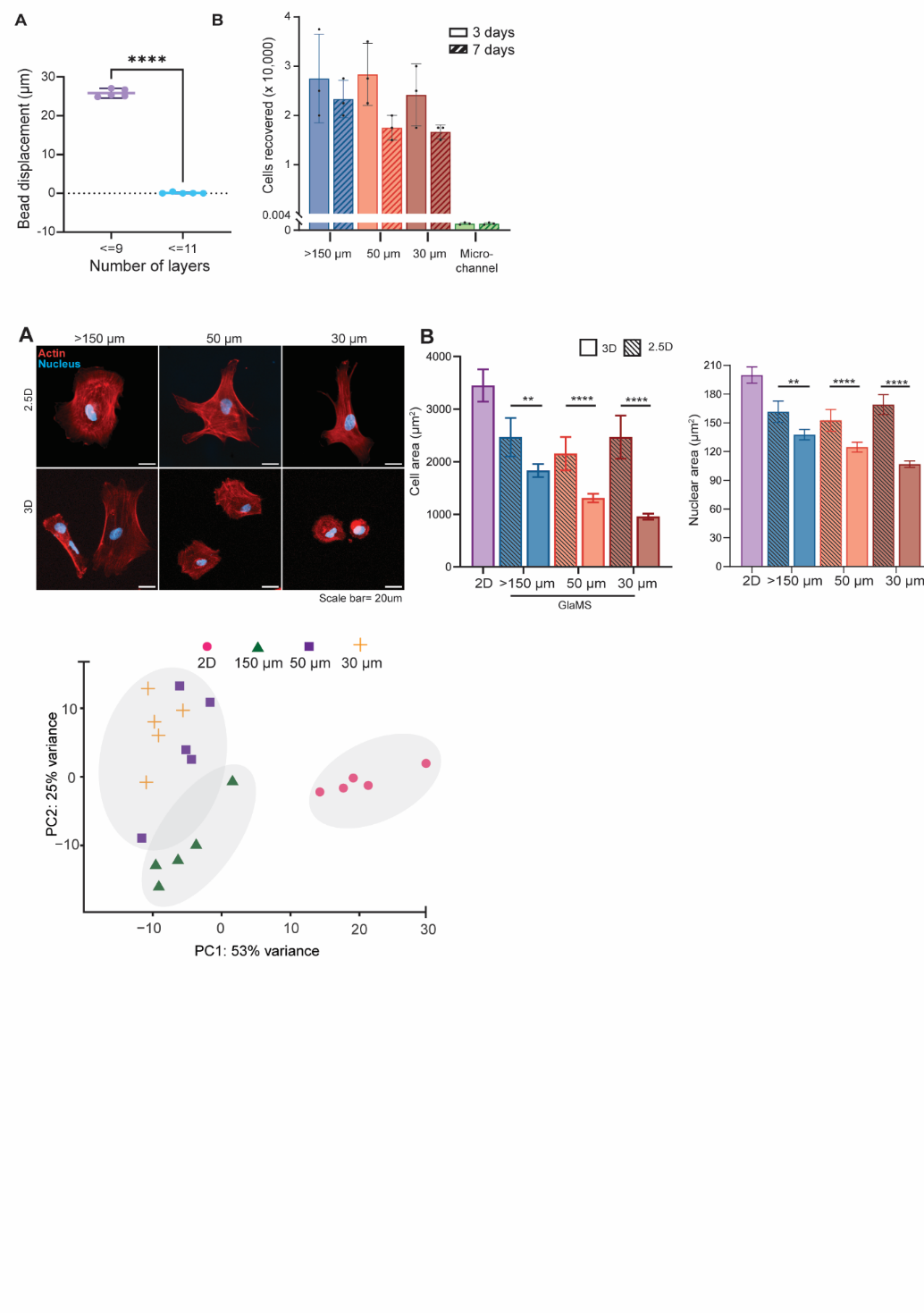

B

***Supplementary Figure 11:*** *A) Methodology used for in-situ lysis of confined cells in GlaMS for RNA sequencing. B) Principal component analysis (PCA) of bulk RNA-seq data.*

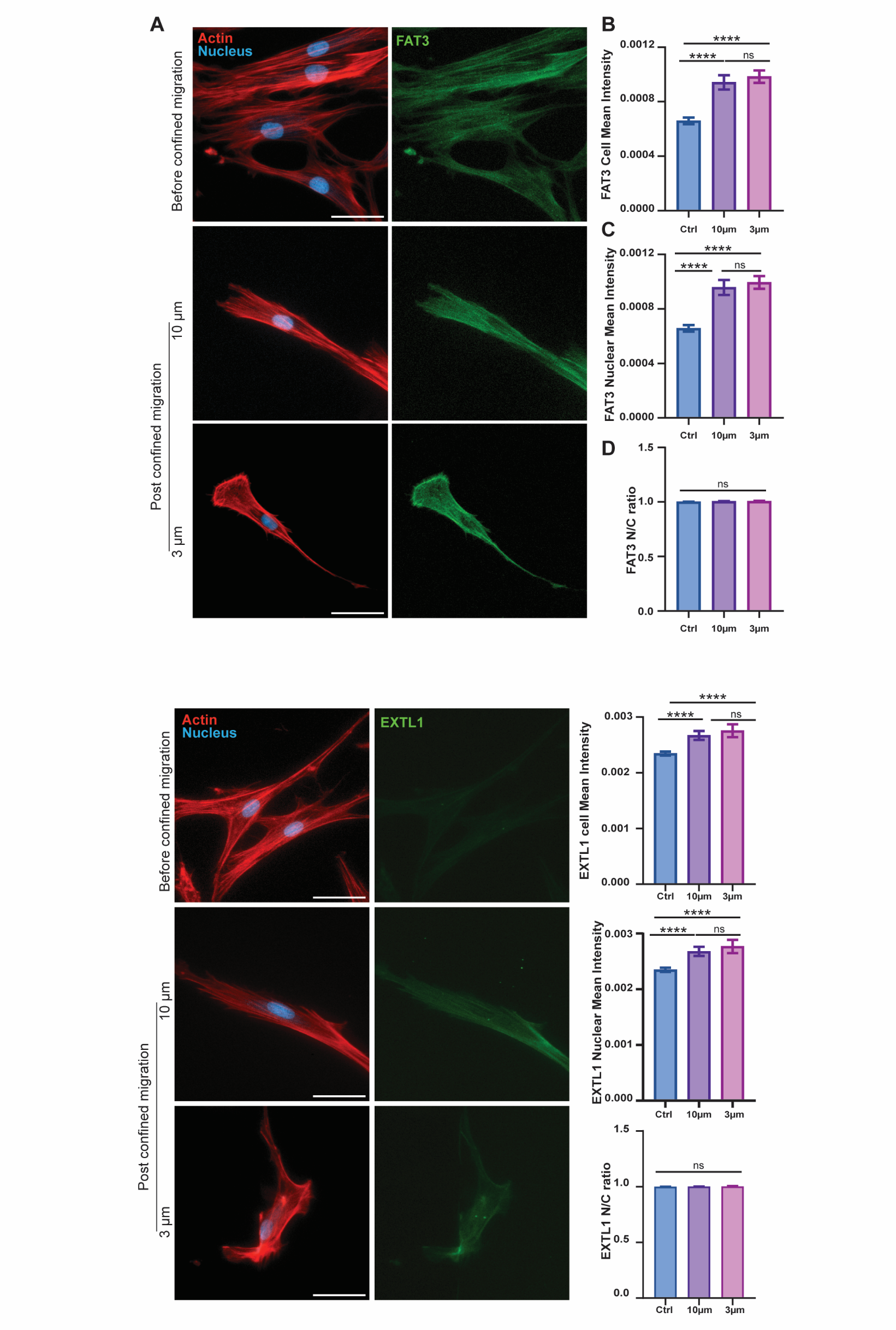

***Supplementary Figure 12: Confinement-sensitive FAT3 regulation.*** *(A) Representative immunofluorescence images of hMSCs before confined migration (Ctrl) and after migration through 10 µm and 3 µm wide channels. Cells are stained for F-actin (red), nuclei (blue), and FAT3 (green). Scale bars: 50 µm. (B) Quantification of FAT3 mean intensity at the whole-cell level. (C) Quantification of nuclear FAT3 mean intensity. (D) Nuclear-to-cytoplasmic (N/C) ratio of FAT3. Graphs represent mean ± 95% confidence interval. Statistical analysis was performed using Kruskal-Wallis test with Dunn’s multiple comparisons post hoc analysis. *P <= 0.05, **P <= 0.01, ***P <= 0.001, ****P <=0.0001.*

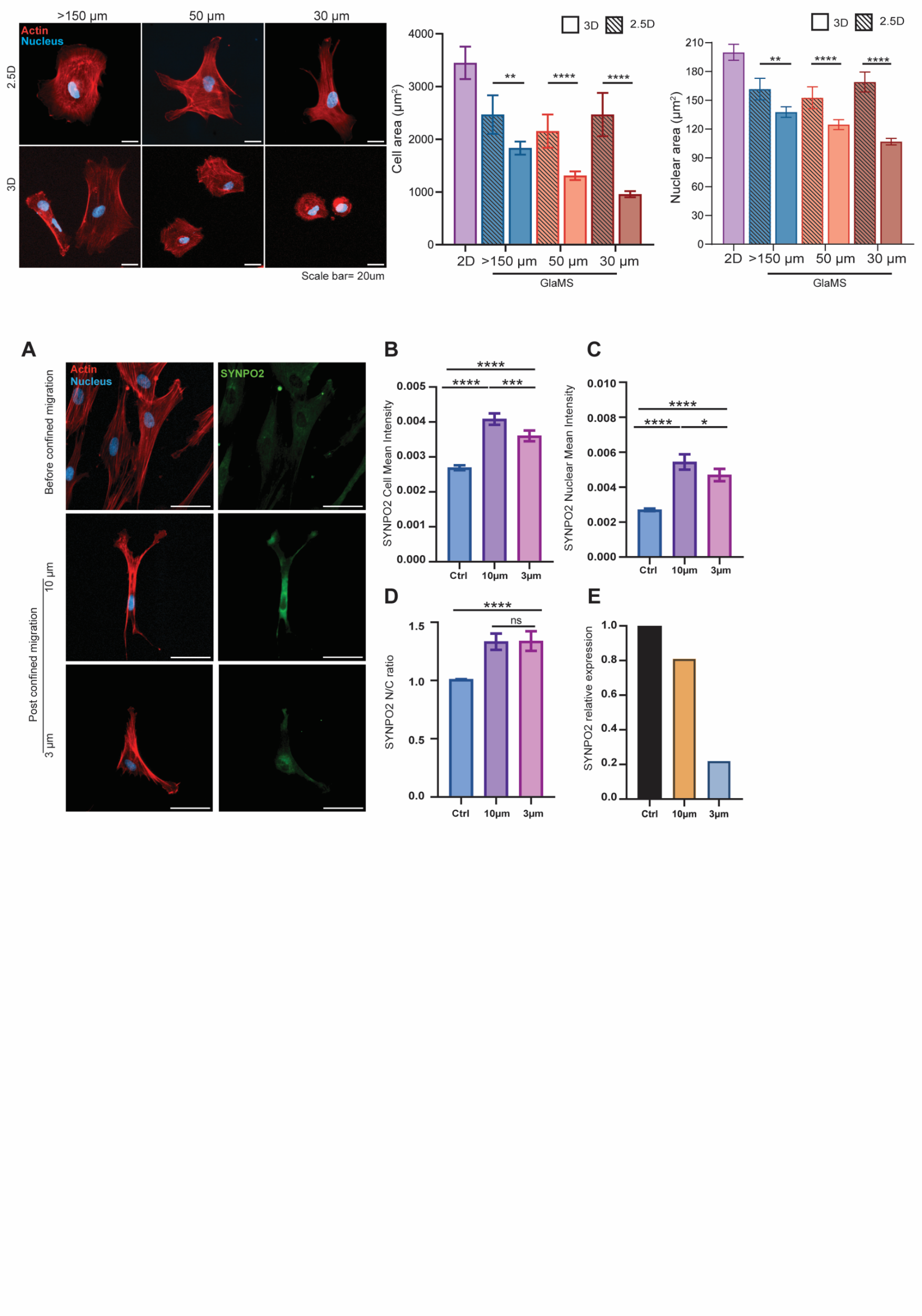

***Supplementary Figure 13: Confinement-sensitive SYNPO2 regulation.*** *(A) Representative immunofluorescence images of hMSCs before confined migration (Ctrl) and after migration through 10 µm and 3 µm wide microchannels. Cells are stained for F-actin (red), nuclei (blue), and SYNPO2 (green). Scale bars: 50 µm. (B) Quantification of SYNPO2 mean intensity at the whole-cell level (C) Quantification of nuclear SYNPO2 mean intensity. (D) Nuclear-to-cytoplasmic (N/C) ratio of SYNPO2.* *Graphs represent mean ± 95% confidence interval. N = 3, n = 72, 53, 40, respectively. (E) Relative expression of SYNPO2 in hMSCs following migration through microchannels (10 µm and 3 µm) compared to unconfined cells, measured by qPCR after cell recovery using Trap-based Recovery After Permeation (TRAP) technique. N = 1, Technical repeats = 3. Statistical analysis was performed using Kruskal - Wallis test with Dunn’s multiple comparisons post hoc analysis. *P <= 0.05, **P <= 0.01, ***P <= 0.001, ****P <=0.0001. Gene expression levels were normalized to GAPDH.*

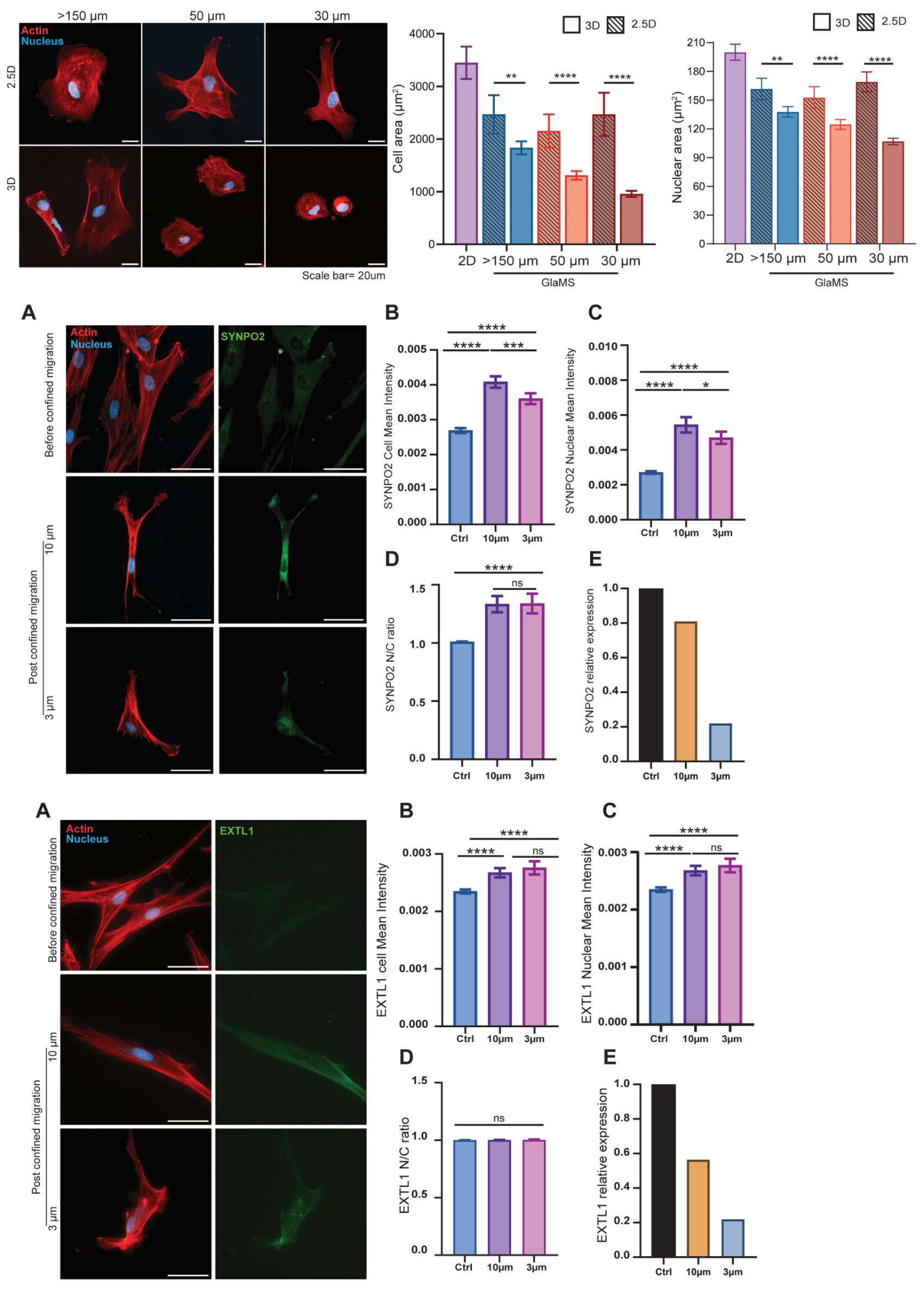

***Supplementary Figure 14: Confinement-sensitive EXTL1 regulation.*** *(A) Representative immunofluorescence images of hMSCs before confined migration (Ctrl) and after migration through 10 µm and 3 µm wide microchannels. Cells are stained for F-actin (red), nuclei (blue), and EXTL1 (green). Scale bars: 50 µm. (B) Quantification of EXTL1 mean intensity at the whole cell level. (C) Quantification of nuclear EXTL1 mean intensity. (D) Nuclear-to-cytoplasmic (N/C) ratio of EXTL1 following confinement.* *Graphs represent mean ± 95% confidence interval. N = 3, n = 85, 52, 41, respectively. (E) Relative expression of EXTL1 in hMSCs following migration through microchannels (10 µm and 3 µm) compared to unconfined cells, measured by qPCR after cell recovery using Trap-based Recovery After Permeation (TRAP) technique. N = 1, Technical repeats = 3. Statistical analysis was performed using Kruskal-Wallis test with Dunn’s multiple comparisons post hoc analysis. *P <= 0.05, **P <= 0.01, ***P <= 0.001, ****P <=0.0001. Gene expression levels were normalized to GAPDH.*

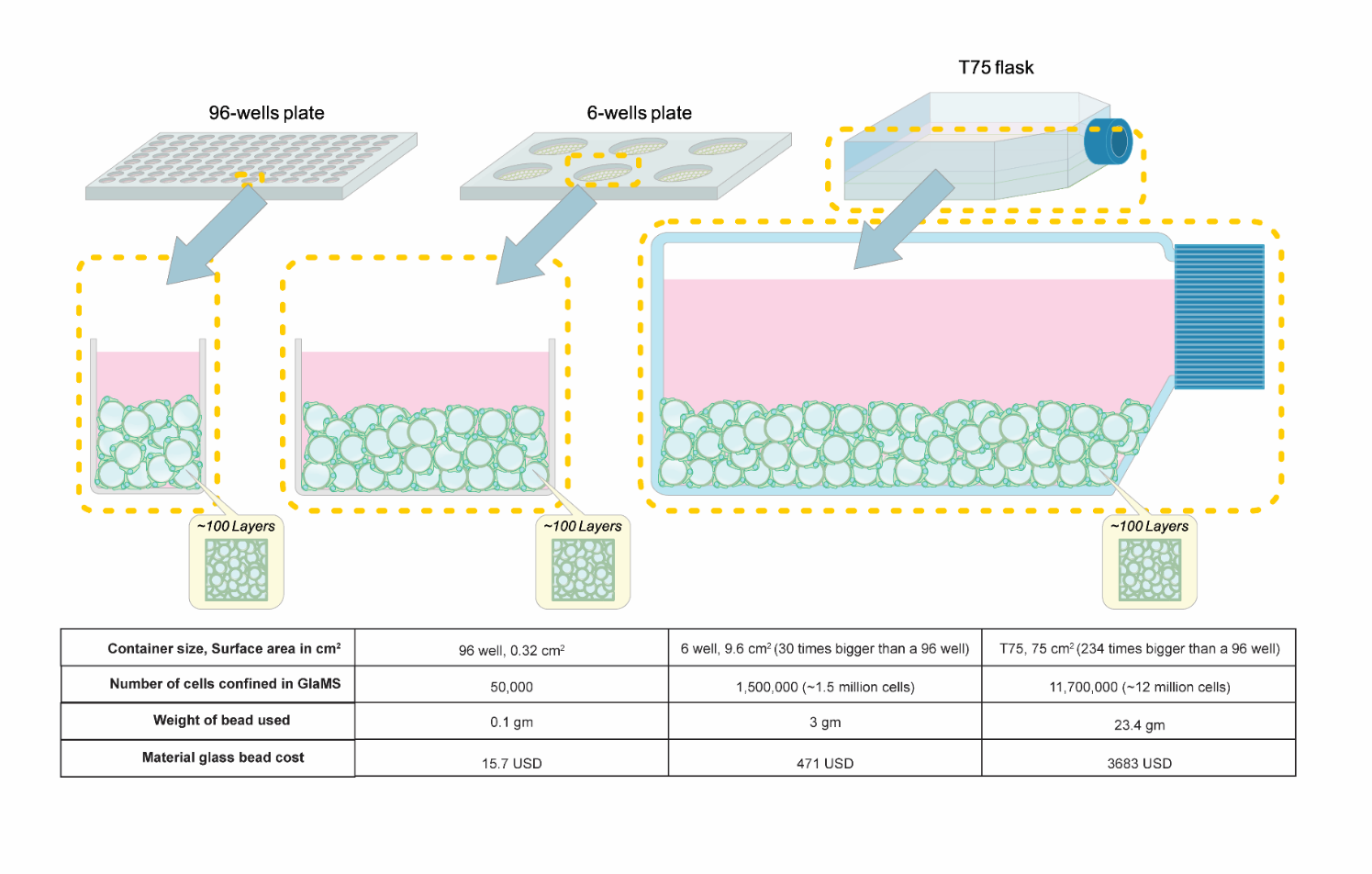

***Supplementary Figure 15: Scaling of the GlaMS platform across different culture plates.*** *Schematic illustrating the scale-up of GlaMS from a single well of a 96-well plate to a 6-well plate and a T-75 flask while maintaining an approximately equivalent scaffold depth of ~100 bead layers.*

**Legends for Supplementary videos:**

***Supplementary video 1:*** *Three-dimensional visualization of interstitial spaces in mouse liver tissue. 3D reconstruction generated using IMARIS from confocal images of mouse liver sections stained for GLUT1 (purple) and nuclei (cyan). The reconstruction highlights the heterogeneous architecture of interstitial spaces within native liver tissue, illustrating the spatial confinement encountered by cells in vivo. Scale bar = 20 µm.*

***Supplementary video 2:*** *Brightfield time-lapse imaging of 30 µm glass microparticles packed in fewer than 9 layers above adherent cells, showing cell-induced displacement and rearrangement of the microparticles. Scale bar = 50 µm.*

***Supplementary Video 3:*** *Brightfield time-lapse imaging of 30 µm glass microparticles packed in more than 11 layers above adherent cells, showing minimal displacement despite cell-generated forces. This demonstrates the stability of the jammed granular scaffold and its ability to maintain consistent confinement geometry. Scale bar = 50 µm.*
